# Construction and application of a technical platform for determining cell cycle- and autophagy-associated cellular accumulation of lipid-based nanoparticles

**DOI:** 10.1101/2024.02.19.579560

**Authors:** Yisha Wang, Gan Luo, Haiyang Wang, Yue Zheng, Xiao Xu, Wenbin Zhou, Junrong Lin, Baocheng Chen, Yifeng Jin, Meihua Sui

## Abstract

Cellular accumulation of biomedical nanoparticles could be affected by cellular biological properties. However, little is known about the influence of cell cycle and autophagy on nanoparticle accumulation. What’s even more tough is that several long-lasting methodological barriers have hampered the experimental performance and restricted related research progress. Herein, a multi-functional platform was constructed for simultaneously overcoming existing obstacles by integrating several technical approaches, particularly mitotic shake-off, for thorough cell cycle phase separation. Strikingly, application of this platform revealed that G2-phase and M-phase cells, two cell populations previously muddled up together as G2/M-phase cells, respectively exhibited the maximum and minimum accumulation of lipid-based nanoparticles. Moreover, although further verification is needed, we have provided a novel line of evidence for enhanced nanoparticle accumulation by autophagy blockade. Besides providing a technical solution, this study discovered characteristic cell cycle- and autophagy-associated nanoparticle accumulations that may offer new insights for optimization and application of nanomedicines.

## Introduction

Leveraging the distinctive surface and size effect, nanodrugs have a wide range of applications in biomedicine^1^. Among a variety of nanocarriers, lipid-based nanoparticles, e.g. liposomes (LIP) and lipid nanoparticles (LNP), serve as exemplary carriers for successful clinical translation^2^. For instance, Doxil^®^, doxorubicin utilizing LIP as the carrier, is the first nano-sized drug approved for patients^3^. BNT162b2, the first LNP-delivered mRNA vaccine, is extensively used worldwide^4^. It has been noted that nanoparticle accumulation could be affected by key cellular biological properties. Cell cycle, a fundamental life process of mammalian cells, is critically involved in the onset, development, treatment response and prognosis of many diseases^5^. Particularly, proliferating tumor cells inside tumor tissues are distributed at distinct cell cycle phases, which is called cell cycle heterogeneity^6,7^. Given that studies from our team and others demonstrate that tumor cells possess cell cycle-dependent sensitivity to a variety of treatments including nano-drugs, cell cycle heterogeneity has been considered as a big challenge for antitumor therapy^6–12^.

Nevertheless, so far only three literatures explored the impact of cell cycle on the retention of inorganic nanoparticles including CdTe QDs^13,14^ and ZnO^15^. Regarding organic nanoparticles, five literatures respectively investigated the cell cycle-associated accumulation of carboxylated polystyrene nanoparticles (PS-COOH), FITC-dextran fibroin-encapsulated microspheres, cRGD-targeting matrix metalloproteinase-sensitive nanoparticles (FITC@NPs-cRGD), folate modified poly (L-amino acid) micelles and Tat/pGL3-Ca^2+^ nanoparticles^16–20^. It is noteworthy that several methodological limitations may interfere with the accuracy of data and result in the discrepant findings previously reported. For instance, the methods used or available in literatures could not differentiate between G2-phase and M-phase cells in physiological state^13–20^. Additionally, cell dissociation agent such as trypsin^13,15–19^, exogenous stimuli (e.g., serum deprivation)^16,18^ and “cell cycle blockers” (e.g., nocodazole)^13,15,18–20^ for artificial synchronization of cell cycle, inevitably disturbed the cellular physiology and subsequent nanoparticle accumulation. Moreover, evaluation of nanoparticle accumulation with confocal laser scanning microscopy (CLSM) was limited at a two-dimensional (2D) level, in which spatial cellular architecture and heterogeneous intracellular distribution of nanoparticles were both ignored^13,17,19,20^. Indeed, these technical barriers have become “bottleneck” problems preventing deeper and further investigations in related research and development.

Autophagy is another major biological process involved in the maintenance of physiological homeostasis and pathogenesis of a wide range of diseases^21^. Interestingly, recent studies indicate that cancer cells are more dependent on autophagy than normal cells, with high levels of endogenous autophagy detected in various types of cancers such as osteosarcoma and pancreatic cancer^22,23^. Evolutionarily conserved autophagy-related genes (*ATG* genes) are core regulatory genes of autophagy^21,24^. Moreover, endosomal/lysosomal system, which is critically involved in nanoparticle accumulation^25^, also plays a pivotal role in autophagy^26^. As a result, there is a growing body of related reports, with the majority focusing on the possible capability of nanoparticles to induce autophagy per se^27^. Up to date four literatures explored the potential effect of autophagy on cellular accumulation of nanoparticles, including NIR-labelled carboxylated polystyrene nanoparticles^28,29^, C6 tagged Ag nanoparticles^30^ and hyaluronic acid-modified MIL-125 nanoparticles loaded with doxorubicin-vitamin E succinate (HA-MIL125@DV) ^31^ all measured at 2D level. Considering that non-specific autophagy inhibitors were used to modulate autophagy in these studies, the obtained data might be confounded by non-autophagy-related pharmacological effects induced by these non-specific inhibitors^32^.

This study was inspired by our strong curiosities on the impact of cell cycle and endogenous autophagy on cellular accumulation of nanoparticles, and by the methodological challenges existed in research on “nano-cell cycle/autophagy” interactions. Herein, a pair of isogenic cancer cell lines including wild type (WT) strain and *ATG7* knockout (*ATG7* KO) derivative was applied to eliminate the interferences caused by non-specific autophagy inhibitors. Importantly, by combining transfection of cell cycle indicator plasmid PIP-FUCCI^33^ and mitotic shake-off^34^, an alive, complete and thorough distinguishment of cells at four distinct cell cycle phases was successfully achieved. Encouragingly, further integration with high-resolution CLSM and three-dimensional (3D) reconstruction technique enabled multi-dimensional evaluations on cellular accumulation of fluorescence-labelled nanoparticles. Through construction and application of this technical platform, cellular accumulation of LIP of Doxil^®^ and LNP of BNT162b2, two widely used nanoparticles, at four distinct cell cycle phases and with specific autophagy status were parallelly and systemically determined respectively. In addition, as a valuable technical solution for simultaneously solving several methodological barriers, the correlations among parameters utilized in the technical platform for determination of nanoparticle accumulation were evaluated.

## Results

### Synthesis and characterization of model nanoparticles DiD-LIP and DiD-LNP

Despite the wide application of LIP and LNP as drug carriers, our understandings on their “nano-bio” interactions are far from sufficient. So far, no literature has investigated the potential impact of either cell cycle or autophagy on the cellular accumulation of LIP or LNP. Hereby LIP and LNP were respectively synthesized according to officially published formulations^35,36^ and our previous report^4^, with DiD as a tracer fluorescence^37,38^. Briefly, DiD-LIP was formulated by hydrating a lipid mixture containing HSPC, cholesterol, mPEG_2000_-DSPE and DiD in sucrose-histidine buffer, followed by downsizing using a 10-mL extruder barrel (Fig. 1a left). DiD-LNP was prepared with a microfluidic mixing device through mixing the ethanol phase containing ALC-0315, DSPC, cholesterol, ALC-159, DiD and the aqueous phase containing firefly luciferase mRNA to enhance the stability of LNP^4,39^ (Fig. 1a right).

Further characterization with dynamic light scattering (DLS) showed that the Z-average/PDI of DiD-LIP and DiD-LNP were 104.433 ± 0.513 nm/0.102 ± 0.036 and 106.767 ± 0.306 nm/0.145 ± 0.009, respectively (Fig. 1b). Additional cryogenic transmission electron microscopy (Cryo-TEM) examination confirmed that DiD-LIP was characterized as hollow spheres, whereas DiD-LNP exhibited a typical electron-dense core structure containing mRNA (Fig. 1c, insert)^4^. Zeta potentials of DiD-LIP and DiD-LNP were -29.100 ± 2.081 mV and 3.660 ± 0.476 mV, respectively (Fig. 1c). These data demonstrate that both model nanoparticles possess highly monodisperse particle-size distribution and weak surface charge.

Importantly, DiD-LIP and DiD-LNP exhibited good stability when stored in phosphate buffer saline (PBS) at 4 °C, without remarkable changes detected for Z-average or PDI from Day 1 to Day 28 (Supplementary Fig. 1a-b; Supplementary Fig. 2), or in DMEM medium containing 10% fetal bovine serum (FBS) at 37 °C for up to 24 hours (simulating the experimental environment) (Fig. 1d-e; Supplementary Fig. 3). Both DiD-LIP and DiD-LNP displayed a single and size-stable intensity peak throughout the whole incubation period and under two distinct physiological conditions. Moreover, the standard curves of phospholipid and corresponding equations were obtained for further dosage determination of DiD-LIP and DiD-LNP (see “**Methods**”, Supplementary Fig. 1c, d)^4^.

**Fig. 1.**
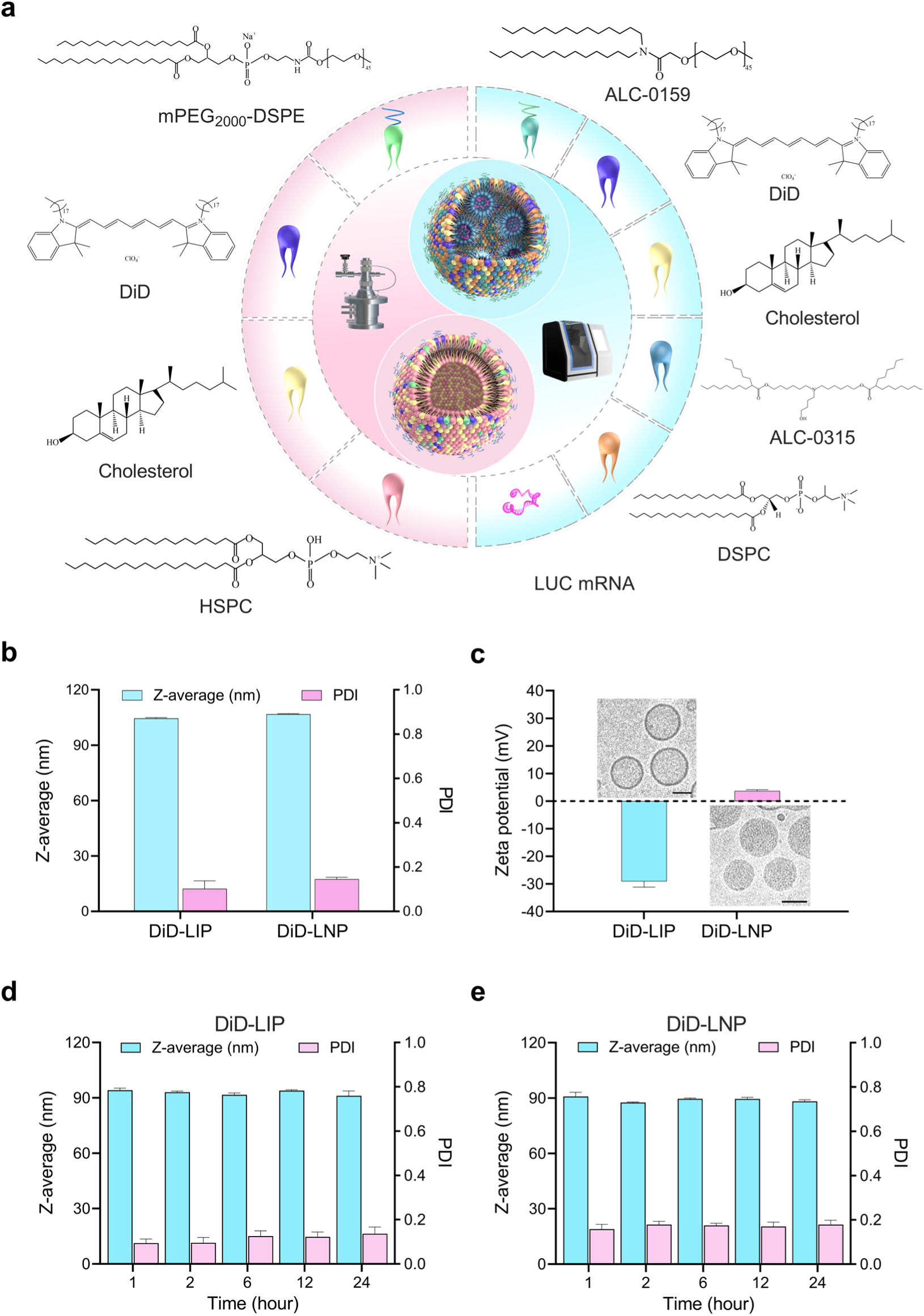
Preparation and characterization of DiD-labelled liposomes (DiD-LIP) and lipid nanoparticles (DiD-LNP). **a**, Schematic illustration of the synthesis of DiD-LIP (left, pink background) and DiD-LNP (right, blue background). Briefly, to synthesis DiD-LIP, the sucrose-histidine buffer hydrated mPEG_2000_-DSPE, DiD, cholesterol and HSPC to generate multilamellar vesicles, followed by size reduction through a 10-mL extruder barrel. To synthesize DiD-LNP, the ethanol phase dissolving ALC-0159, DiD, cholesterol, ALC-0315 and DSPC was mixed with the aqueous phase containing firefly luciferase mRNA (LUC mRNA) at a flow rate ratio of 1: 3 (ethanol: aqueous) via a microfluidic mixing device. **b-c**, Z-average, polydispersity index (PDI), Zeta potentials and representative Cryo-TEM images (inserts) of DiD-LIP and DiD-LNP. Scale bars, 50 nm. **d-e**, Z-average and PDI of DiD-LIP and DiD-LNP in DMEM containing 10% FBS stored at 37 °C for up to 24 hours. Data in **b-e** were presented as “mean ± standard deviation” of three independent experiments. Z-average and PDI among various time points were evaluated by ANOVA with Bonferroni correction, with no statistically significant difference detected (all *P* > 0.05).

### Establishment of a multi-functional technical platform for determining cell cycle- and autophagy-associated cellular accumulation of nanoparticles

Using empty vector as a control, the isogenic WT and *ATG7* KO U2OS cell lines previously constructed were transduced with lentiviral particles carrying the cell cycle-indicating PIP-FUCCI plasmid (see “**Methods**”). After screening with G418 and flow cytometry, corresponding WT and *ATG7* KO PIP-FUCCI cell lines expressing strong mVenus (green fluorescence) and/or mCherry (red fluorescence) were obtained (Fig. 2a). PIP-FUCCI transfection and subsequent selection had marginal influence on cellular morphology and proliferation rate (Supplementary Figs. 4 and 5).

However, although single positivity of mVenus and mCherry could respectively indicate cells at G1-phase and S-phase, both G2-phase and M-phase cells are mVenus and mCherry double-positive and exhibit yellow fluorescence (overlap of green and red). That is, PIP-FUCCI transfection by itself cannot distinguish between G2-phase and M-phase cells in situ. Fortunately, different from G1-phase, S-phase and G2-phase cells naturally adhering to the culture dish with polyhedral shapes, typical M-phase cells have unique properties such as spherical shape^40^ and significantly reduced adhesion capability or suspended state in culture media due to “mitotic rounding”, leading to convenient collection of M-phase cells through gentle mitotic shake-off^34^. Enlightened by these characteristics, we combined PIP-FUCCI transfection with mitotic shake-off, through which an approach thoroughly distinguishing four cell cycle phases was established. In addition, labelling cell membrane with FITC-EpCAM prior to mitotic shake-off helped to exclude apoptotic cells and cellular debris via drawing the outline of cell membrane (see “**Methods**”; Fig. 2b, highlighted with dashed frame).

Further integration of high-resolution CLSM and 3D reconstruction technique significantly strengthened this multi-functional platform for determining cell cycle- and autophagy-associated cellular accumulation of nanoparticles (Fig. 2b). Using DiD-LIP (blue fluorescence) as a representative, nanoparticle accumulation was not only characterized at different dimensional levels, but respectively evaluated at four distinct cell cycle phases in WT and *ATG7* KO PIP-FUCCI cells (Fig. 2c). In brief, this new platform has multiple superiorities including complete and thorough separation and collection of alive cells at four cell cycle phases, elimination of interferences caused by “extra” treatments (e.g. cell dissociation agents, cell cycle synchronization drugs, none-specific autophagy inhibitors), and capability for parallel investigations on multiple biological parameters/dimensions.

**Fig. 2.**
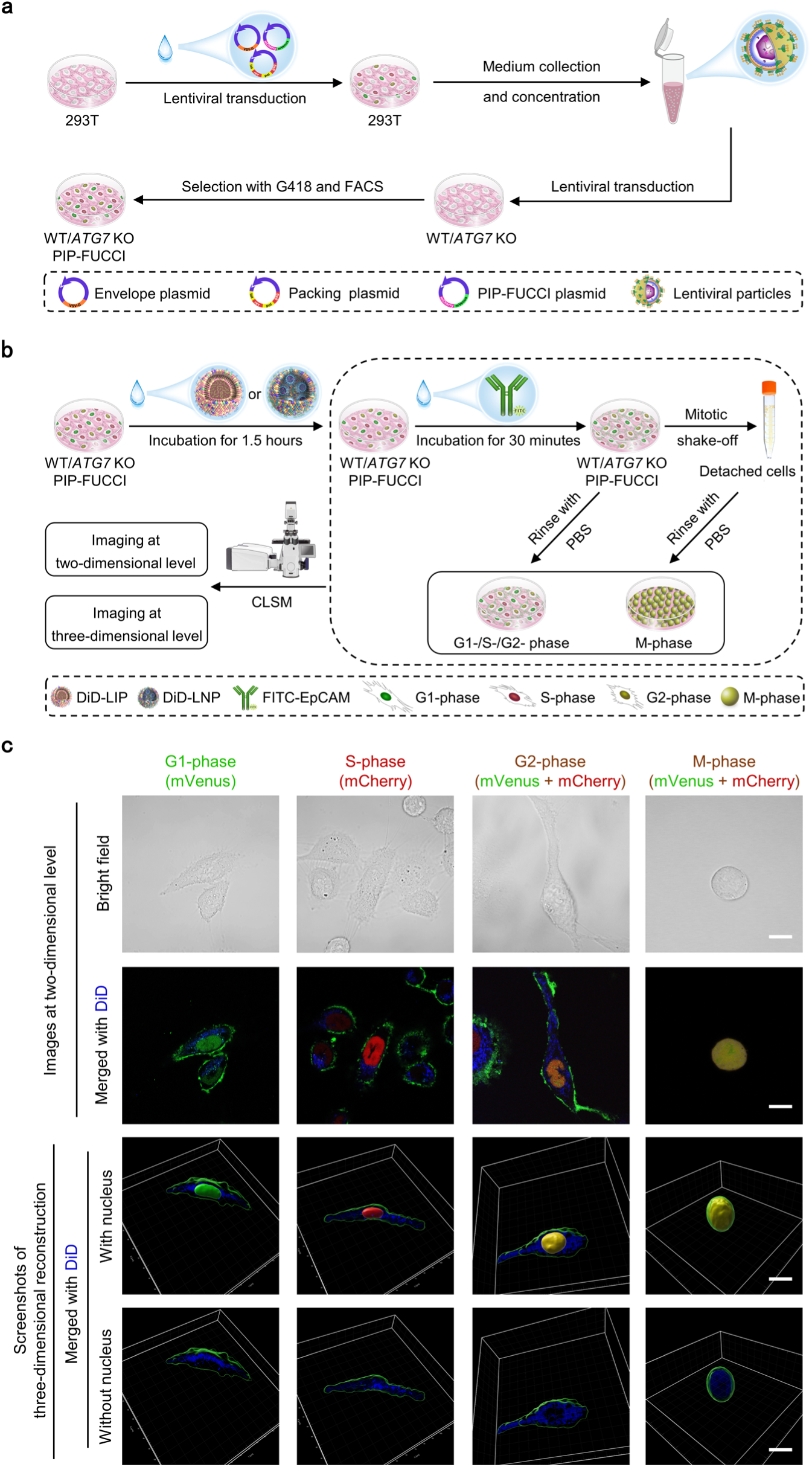
Establishment of an alive and multi-functional technical platform for determining cell cycle- and autophagy-associated nanoparticle accumulation. **a,** Schematic diagram shows the construction of cell lines stably expressing PIP-FUCCI (cell cycle indicator) plasmid. Envelope plasmid PMD2.G, packing plasmid psPAX2 and transfer plasmid PIP-FUCCI were simultaneously added into 293T cells. Afterwards, the culture medium was collected and concentrated, followed by application of lentiviral supernatant containing PIP-FUCCI to isogenic wide type (WT) and *ATG7* knockout (*ATG7* KO) U2OS cell lines. Subsequently, WT and *ATG7* KO PIP-FUCCI cell lines stably expressing PIP-FUCCI were obtained after selection with G418 and flow cytometry (FACS). **b**, Schematic diagram indicates the establishment of a multi-functional platform for determining cell cycle phase- and autophagy-associated nanoparticle accumulation. WT and *ATG7* KO PIP-FUCCI cells were treated with DiD-labelled liposomes (DiD-LIP, equivalent to 5 μg/mL DOX) or lipid nanoparticles (DiD-LNP, equivalent to 6 μg/mL luciferase mRNA) for 1.5 hours, followed by FITC-EpCAM incubation for 30 minutes to label cell membrane (green cell membrane). Complete distinguishment and collection of cells at four distinct cell cycle phases was achieved through combining PIP-FUCCI expression to indicate G1-phase (mVenus positive, green nucleus), S-phase (mCherry positive, red nucleus) and G2-phase (mVenus and mCherry double-positive, yellow nucleus) cells attached to culture dish, with mitotic shake-off to collect detached or floating M-phase cells (mVenus and mCherry double-positive, yellow nucleus). Further integration of confocal laser scanning microscope (CLSM) and three-dimensional reconstruction technique enabled detection and calculation of nanoparticle accumulation at both two-dimensional and three-dimensional levels. **c**, Representative light microscopic and CLSM images demonstrating the successful construction and application of new technical platform, using DiD-LIP (blue fluorescence) and WT PIP-FUCCI cells as a model system. Scale bars, 20 μm.

It is noteworthy that although cells harvested with mitotic shake-off have been described as all M-phase cells^41^, a rare population of just divided spherical daughter cells having not fully re-adhered to the dish might be simultaneously collected. However, daughter cells just completed mitosis have actually entered a new cell cycle and are at G1-phase. Due to rapid decay of mCherry protein, daughter cells quickly become mVenus single-positive displaying green nuclear fluorescence, which differs from the pre-division mitotic cells with yellow nuclear fluorescence^33^. Therefore, suspended daughter cells will neither be selected nor analyzed as M-phase cells by mistake in this study.

To further confirm these characteristics, routinely cultured WT and *ATG7* KO PIP-FUCCI cells without exposure to nanoparticles were monitored with CLSM for up to 25 hours (estimated cell doubling time, Supplementary Fig. 6). Using individually tracked cancer cells as examples, our “longitudinal” data further demonstrated the reliability of combining PIP-FUCCI expression with morphological examination for precise cell cycle indication throughout a complete cell cycle including “M-to-G1” phase transition (Supplementary Fig. 6). Moreover, our findings indicating gradual increases in cell volume while cell cycle progression have provided another piece of evidence that the M-phase cells analyzed herein were pre-division mother cells with maximum cell volumes, but not just divided daughter cells minimum cell volumes (see subsequent “**Results**”).

### Little influence of DiD-LIP and DiD-LNP per se on cell cycle distribution and autophagy of WT and *ATG7* KO PIP-FUCCI cell lines

To clarify the impact of cell cycle and autophagy on nanoparticle accumulation, the possible interference of model nanoparticles on cancer cells per se needs to be firstly excluded. Therefore, WT and *ATG7* KO PIP-FUCCI cells were respectively treated with DiD-LIP and DiD-LNP for further evaluation. As depicted in Fig. 3a and Supplementary Table 1, flow cytometry analysis showed that percentages of nanoparticle-untreated cancer cells at G1-phase, S-phase, G2-phase and M-phase were 33.10 ± 1.77%, 54.83 ± 2.82%, 11.81 ± 0.93% and 0.26 ± 0.19%, respectively, for WT PIP-FUCCI cell line, and 31.30 ± 2.21%, 54.22 ± 2.49%, 13.94 ± 1.29% and 0.53 ± 0.16%, respectively, for *ATG7* KO PIP-FUCCI cell line. Percentages of DiD-LIP-treated WT and *ATG7* KO PIP-FUCCI cell lines at G1-phase, S-phase, G2-phase and M-phase were 33.18 ± 2.08%, 54.64 ± 3.14%, 11.92 ± 1.09% and 0.26 ± 0.17%, respectively, and 31.25 ± 2.15%, 53.99 ± 2.37%, 14.29 ± 0.57%, 0.48 ± 0.10%, respectively. Correspondingly, the cell cycle distributions of DiD-LNP-treated WT and *ATG7* KO PIP-FUCCI cell lines at G1-phase, S-phase, G2-phase and M-phase were 33.70 ± 2.36%, 54.38 ± 3.37%, 11.66 ± 0.92% and 0.27 ± 0.21%, respectively, and 31.15 ± 0.61%, 55.02 ± 1.58%, 13.27 ± 1.44% and 0.56 ± 0.16%, respectively. Importantly, further statistical analysis demonstrated that no matter with or without nanoparticle treatment, no significant difference on cell cycle distribution was observed in WT and *ATG7* KO PIP-FUCCI cells (Supplementary Table 2).

Moreover, data obtained with additional regular flow cytometry assay (without mitotic shake-off) at longer time points demonstrated that in both WT and *ATG7* KO PIP-FUCCI cell lines, neither DiD-LIP nor DiD-LNP exhibited significant influence on general cell cycle distribution after administration for 12 hours, 24 hours and 48 hours, respectively (Supplementary Figs. 7 and 8, Supplementary Table 3 and Table 4).

**Fig. 3.**
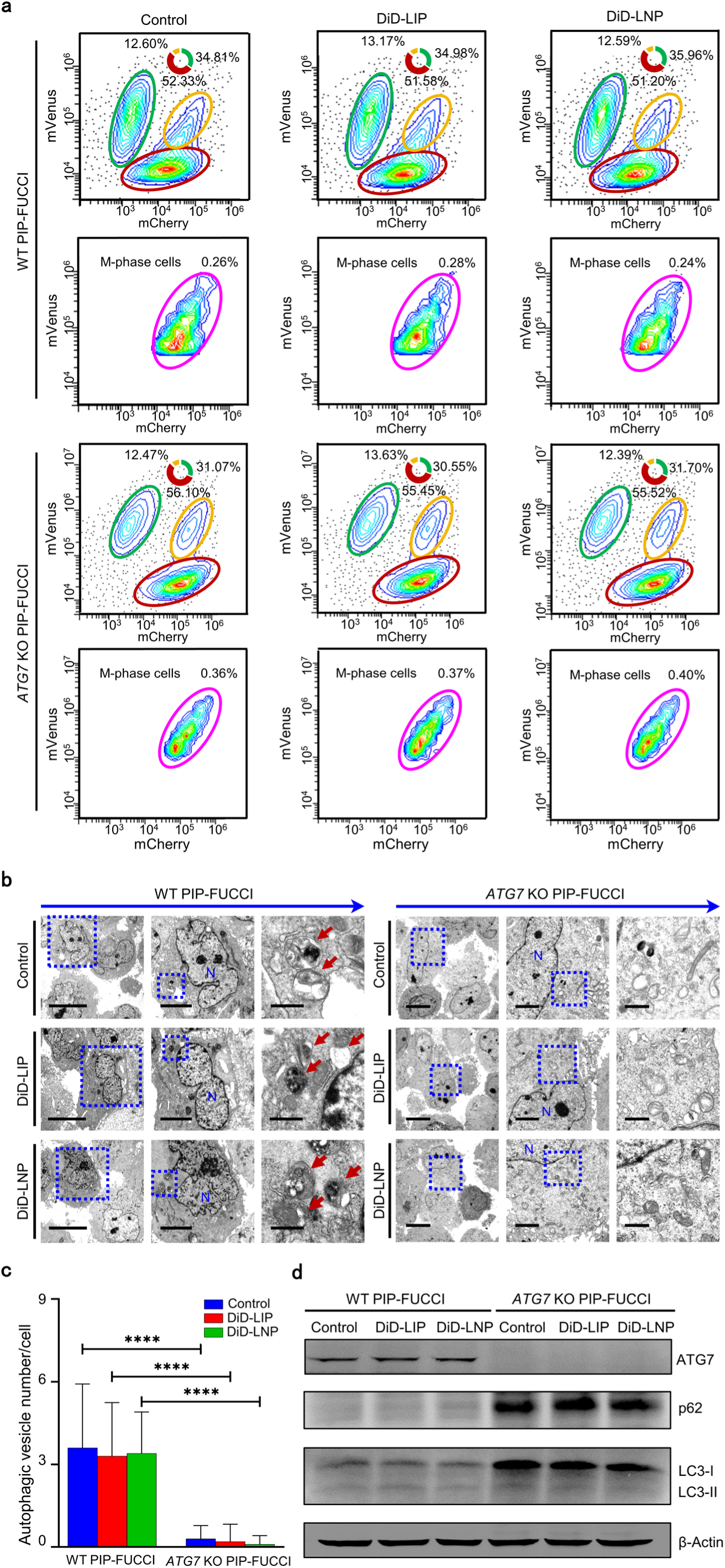
Validation of cell cycle distribution and endogenous autophagy in WT and *ATG7* KO PIP-FUCCI cells with or without nanoparticle treatment. **a**, Representative flow cytometry data on thorough cell cycle distribution. Cells accumulated at G1-phase, S-phase and G2-phase (upper panel for each cell line), as well as M-phase (isolated with mitotic shake-off; lower panel for each cell line) were respectively circled in green, red, orange and purple, with corresponding percentages indicated. **b**, Representative transmission electron microscope images of autophagic vesicles indicated by rust-red arrows. N, nucleus. Scale bars, from left to right, 10 μm, 4 μm and 1 μm for WT PIP-FUCCI cells, and 10 μm, 3 μm and 1 μm for *ATG7* KO PIP-FUCCI cells. **c**, Statistical analysis of the autophagic vesicle number per cell. Data were presented as “mean ± standard deviation” (n = 10) and further analyzed by Mann-Whitney U test. ****, *P* < 0.0001. **d**, Representative western blot analysis of autophagy-related proteins including ATG7, p62 and LC3. WT PIP-FUCCI, wide type U2OS cells stably expressing PIP-FUCCI; *ATG7* KO PIP-FUCCI, *ATG7* knockout U2OS cells stably expressing PIP-FUCCI. DiD-labelled liposomes (DiD-LIP) and lipid nanoparticles (DiD-LNP) were administered to cancer cells at a dose equivalent to 5 μg/mL DOX and a dose equivalent to 6 μg/mL luciferase mRNA, respectively for 2 hours.

Previous studies demonstrate that *ATG7* knockdown effectively blocked cellular autophagy^42^. Indeed, although autophagic vesicles (AVs, including autophagosomes and autophagic lysosomes) with typical double-layer membrane structure were easily observed in WT PIP-FUCCI cells with TEM, no typical AVs were seen in *ATG7* KO PIP-FUCCI cells (Fig. 3b). Further statistical analysis confirmed significantly fewer AVs in *ATG7* KO PIP-FUCCI cells than in WT PIP-FUCCI cells (Fig. 3c; Supplementary Table 5). As anticipated, ATG7 protein level was high in WT PIP-FUCCI cells while undetectable in *ATG7* KO PIP-FUCCI cells in Western blotting (Fig. 3d). Moreover, LC3-II/LC3-I ratio, which is positively correlated with the autophagy level^43^, was significantly decreased in *ATG7* KO PIP-FUCCI cells. In contrast, p62, which is negatively correlated with autophagy^43^, was dramatically increased in *ATG7* KO PIP-FUCCI cells. Our data firmly demonstrated the successful blockade of autophagy in *ATG7* KO PIP-FUCCI cells. Encouragingly, neither DiD-LIP nor DiD-LNP exhibited any effect in aforementioned assays, indicating that designated nanoparticle treatment had marginal impact on the autophagy of WT and *ATG7* KO PIP-FUCCI cells.

### Determination of cell cycle- and endogenous autophagy-associated cellular accumulation of DiD-LIP

Next, we meticulously captured abundant CLSM images with intracellular co-localization of cell cycle-indicating fluorescence and DiD-LIP using this new platform (Supplementary Videos 1-8; Supplementary Fig. 9). Subsequently, cell size, Total Fluorescence Intensity (TFI) and Mean Fluorescence Intensity (MFI) were respectively calculated and analyzed at 2D and 3D levels (Fig. 4; Table 1; Supplementary Tables 6 and 7). Our data showed that TFI (2D) of DiD-LIP across various cell cycle phases exhibited same characteristics in both cell lines, which was M < G1 ≈ S < G2. Further integration of 3D reconstruction technique revealed the following cell cycle-associated characteristics of TFI (3D) of DiD-LIP: M (≈ G1) < (G1 ≈) S < G2 in WT PIP-FUCCI cells; M < G1 < S < G2 in *ATG7* KO PIP-FUCCI cells.

**Table 1.**
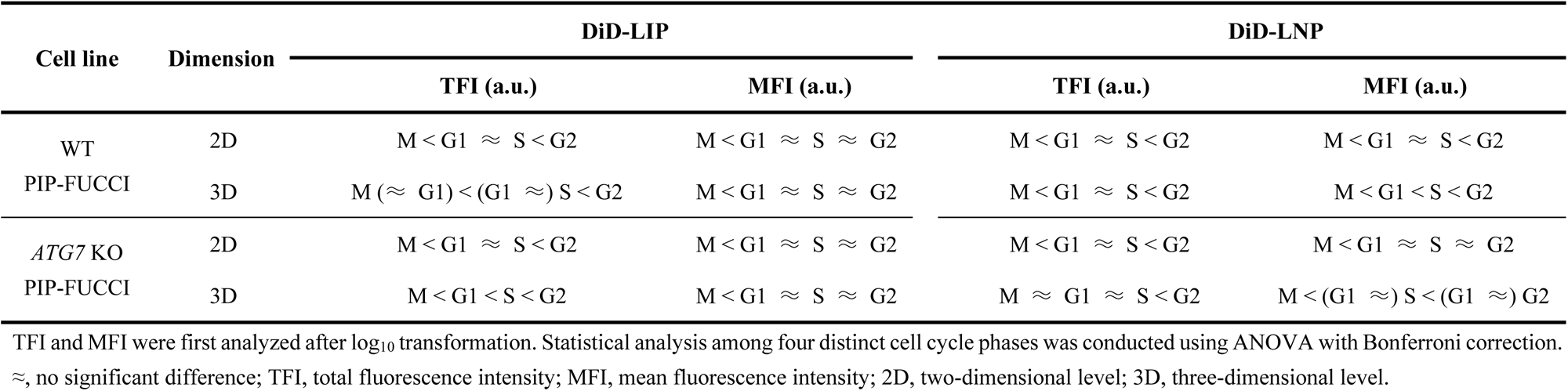
Cell cycle- and autophagy-associated accumulation patterns of DiD-LIP and DiD-LNP by cancer cells based on the multi-functional platform.

As expected, both cell lines exhibited a gradual increase in cell area (2D level) upon cell cycle progressing from G1-phase to G2-phase, followed by a decrease at M-phase due to “mitotic rounding” to become spherical^44^. Consistent with previous reports, cell volume (3D level) gradually increased while cells sequentially going through G1-phase, S-phase, G2-phase and M-phase of the cell cycle^45,46^. Furthermore, MFI (2D) based on cell area and MFI (3D) based on cell volume were initially calculated and analyzed. Impressively, MFI (2D) and MFI (3D) of DiD-LIP exhibited same cell cycle-associated characteristics in both cell lines, which was M < G1 ≈ S ≈ G2. These data may suggest that the relatively higher increase in cell size of G2-phase cells attenuated the aforementioned difference in TFI detected between G2-phase and G1-phase/S-phase. Taken together, our study not only differentiated between G2-phase and M-phase cells in physiological state, but provided initial evidence that cells at G2-phase and M-phase respectively had the maximum and minimum nanoparticle accumulation. Interestingly, it has been suggested that M-phase cells have declined endocytosis and elevated cell membrane tension, which may assist mother cell to prevent external disturbances during mitosis^47^. These properties support our findings that M-phase cells had minimum nanoparticle accumulation among four cell cycle phases.

As mentioned above, discrepant results were reported in literatures investigating the influence of cell cycle on nanoparticle accumulation^13–20^. For instance, although accumulation of CdTe QDs^13,14^, ZnO^15^, PS-CCOH^16^ and FITC–dextran fibroin-encapsulated microspheres^17^ showed same cell cycle-associated characteristics (G2/M > S > G1), FITC@NPs-cRGD accumulation exhibited a reverse ranking (G1 > S > G2/M). It should be noted that cells at G2-phase and M-phase were muddled up as G2/M-phase cells in most previous studies^13–19^, except G2-phase and M-phase synchronizations were respectively induced using RO3306 and paclitaxel in a study investigating cell-cycle-dependent macropinocytosis of Tat/pDNA-Ca^2+^ nanoparticles^20^.

**Fig. 4.**
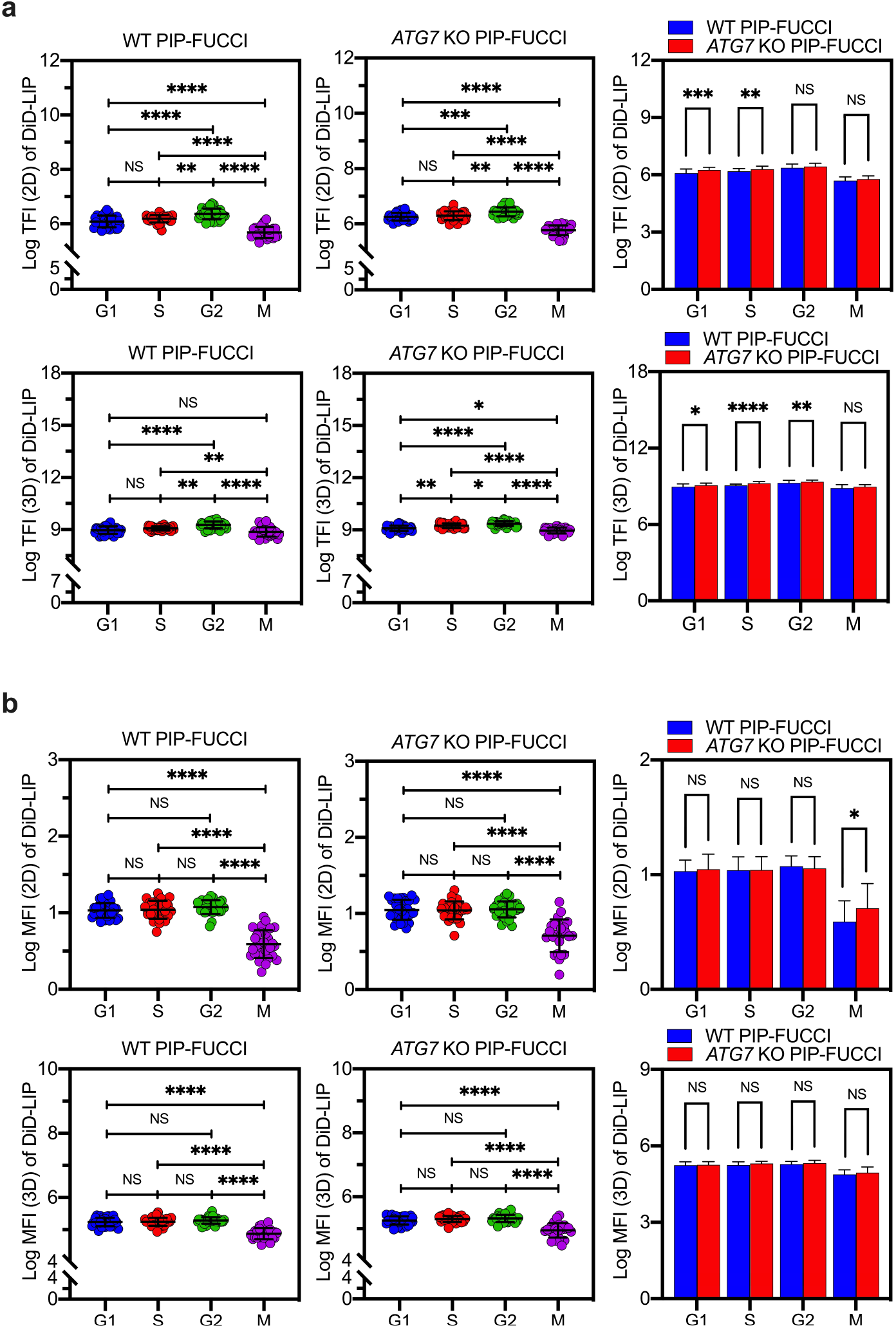
Determination of cell cycle- and endogenous autophagy-associated cellular accumulation of DiD-labelled liposomes (DiD-LIP) using the newly established technical platform. **a**, Statistical analysis of Total Fluorescence Intensity (TFI) in G1-phase, S-phase, G2-phase and M-phase WT and *ATG7* KO PIP-FUCCI cells respectively measured at two-dimensional (2D, upper panel) and three-dimensional (3D, lower panel) levels. **b**, Statistical analysis of Mean Fluorescence Intensity (MFI) in WT and *ATG7* KO PIP-FUCCI cells respectively measured at aforementioned four cell cycle phases, as well as at 2D (upper panel) and 3D (lower panel) levels. WT PIP-FUCCI, wide type U2OS cells stably expressing PIP-FUCCI; *ATG7* KO PIP-FUCCI, *ATG7* knockout U2OS cells stably expressing PIP-FUCCI. DiD-LIP was administered at a dose equivalent to 5 μg/mL DOX for 2 hours. All data were analyzed after log_10_ transformation and presented as “mean ± standard deviation” (n = 25-30). Statistical analysis of DiD-LIP accumulation among four distinct cell cycle phases was conducted using ANOVA with Bonferroni correction, while differences between WT and *ATG7* KO PIP-FUCCI cell lines were evaluated with an independent sample *t*-test. *, *P* < 0.05; **, *P* < 0.01; ***, *P* < 0.001; ****, *P* < 0.0001. NS, not significant.

Moreover, both TFI and MFI of DiD-LIP suggested an association between nanoparticle accumulation and endogenous autophagy. For instance, TFI (2D) of G1-phase and S-phase *ATG7* KO PIP-FUCCI cells, TFI (3D) of G1-phase, S-phase and G2-phase *ATG7* KO PIP-FUCCI cells, as well as MFI (2D) of M-phase *ATG7* KO PIP-FUCCI cells, all exhibited significant increases and surpassed those of WT PIP-FUCCI cells. Previously there are four literatures explored the influence of autophagy on nanoparticle accumulation, all of which using 2D fluorescence intensity for accumulation assessment and non-specific autophagy inhibitors for autophagy inhibition. Among these reports, Fageria et al. reported a reduced cellular accumulation of Ag nanoparticles upon chloroquine treatment^30^, and Sipos et al. observed a suppressed cellular accumulation of polystyrene nanoparticles after exposure to 3-methyladenine and bafilomycin A1^28,29^. However, interestingly, co-loading of 3-methyladenine into HA-MIL125@DV nanoparticles (so called HA-MIL125@DVMA) led to significantly increased nanoparticle accumulation in both drug-sensitive MCF-7 and multidrug-resistant MCF-7/ADR cell lines^31^. Although multiple contributing factors such as nanoparticle- and/or cell line-specificity might be involved in mediating the different findings, it is noteworthy that different from *ATG7* knockout that specifically blocks endogenous autophagy, non-autophagy-related pharmacological effects such as induction of cell death and lysosomotropic activities have been reported for chloroquine, 3-methyladenine and bafilomycin A1, which may interfere with the cellular accumulation of nanoparticles^48,49^.

### Determination of cell cycle- and endogenous autophagy-associated cellular accumulation of DiD-LNP

WT and *ATG7* KO PIP-FUCCI cells were further treated with DiD-LNP and evaluated with the multi-functional platform (Fig. 5; Table 1; Supplementary Videos 9-16; Supplementary Fig. 10; Supplementary Tables 8 and 9). Impressively, TFI (2D) of DiD-LNP in both cell lines, and TFI (3D) and MFI (2D) of DiD-LNP in WT PIP-FUCCI cell line, all exhibited same cell cycle-associated characteristics as TFI (2D) of DiD-LIP, which was M < G1 ≈ S < G2. MFI (3D) in WT and TFI (3D) in *ATG7* KO PIP-FUCCI cells showed similar characteristics, except G1 < S in WT and M ≈ G1 in *ATG7* KO PIP-FUCCI cells. Moreover, MFI (2D) and MFI (3D) in *ATG7* KO PIP-FUCCI cells were M < G1 ≈ S ≈ G2 and M < (G1 ≈) S < (G1 ≈) G2, respectively, coinciding with the aforementioned attenuation of differences between G2-phase and G1-phase/S-phase due to relatively higher increase in cell size of G2-phase cells. In general, multi-dimensional data obtained with DiD-LNP confirmed that G2-phase cells had maximum whereas M-phase cells had minimum nanoparticle accumulation. Interestingly, we noted that mean values of TFI and MFI of two model nanoparticles exhibited similar tendency in both cell lines, which is M < G1 < S < G2. These data might imply that when proliferating cells progress into a new cell cycle with physiological activities such as cellular protein and DNA synthesis gradually intensifying, nanoparticle accumulation becomes active in synchronization till the cells enter “torpid” M-phase^50,51^.

Moreover, it is noteworthy that M-phase cells have the minimum cell areas, which would weaken the difference of MFI (2D) between M-phase cells and cells at other phases. Nevertheless, M-phase cells of both cell lines still exhibited extremely significant decrease in MFI (2D) upon treatment with two model nanoparticles (nearly all *P* < 0.0001). In contrast, M-phase cells have the maximum cell volumes, which would strengthen the difference of MFI (3D) between M-phase cells and cells at other phases. It is thereby logical that the following significant differences in MFI (3D) arose, although no difference was detected with corresponding TFI (3D): M < G1 in MFI (3D) of DiD-LIP in WT PIP-FUCCI cells, M < G1 and M < S in MFI (3D) of DiD-LNP in *ATG7* KO PIP-FUCCI cells, all *P* < 0.0001.

Interestingly, different from DiD-LIP that exhibited several significantly higher TFI and MFI in *ATG7* KO PIP-FUCCI cells than those in WT PIP-FUCCI cells, DiD-LNP showed increased TFI but reduced MFI in *ATG7* KO PIP-FUCCI cells compared with WT strain. In detail, TFI (2D) of M-phase, and TFI (3D) of G1-phase and M-phase *ATG7* KO PIP-FUCCI cells were significantly higher than those of WT PIP-FUCCI cells. However, both MFI (2D) and MFI (3D) of S-phase and G2-phase *ATG7* KO PIP-FUCCI cells were significantly lower than those in WT strain. These data indicate that autophagy might have diverse influences on cellular accumulation of nanoparticles which deserves further investigation. Moreover, although cellular accumulation of DiD-LIP and DiD-LNP exhibited several common characteristics, varied features were also observed between these two nanoparticles regarding cell cycle- and autophagy-associated cellular accumulation. These variations might attribute to different physiochemical properties between DiD-LIP and DiD-LNP, e.g. surface charge, nanoparticle composition and spatial structure, leading to distinctive “nano-bio” interactions such as different characteristics in endocytosis, subsequent endo-lysosomal escape, degradation and even exocytosis of nanoparticles.

**Fig. 5.**
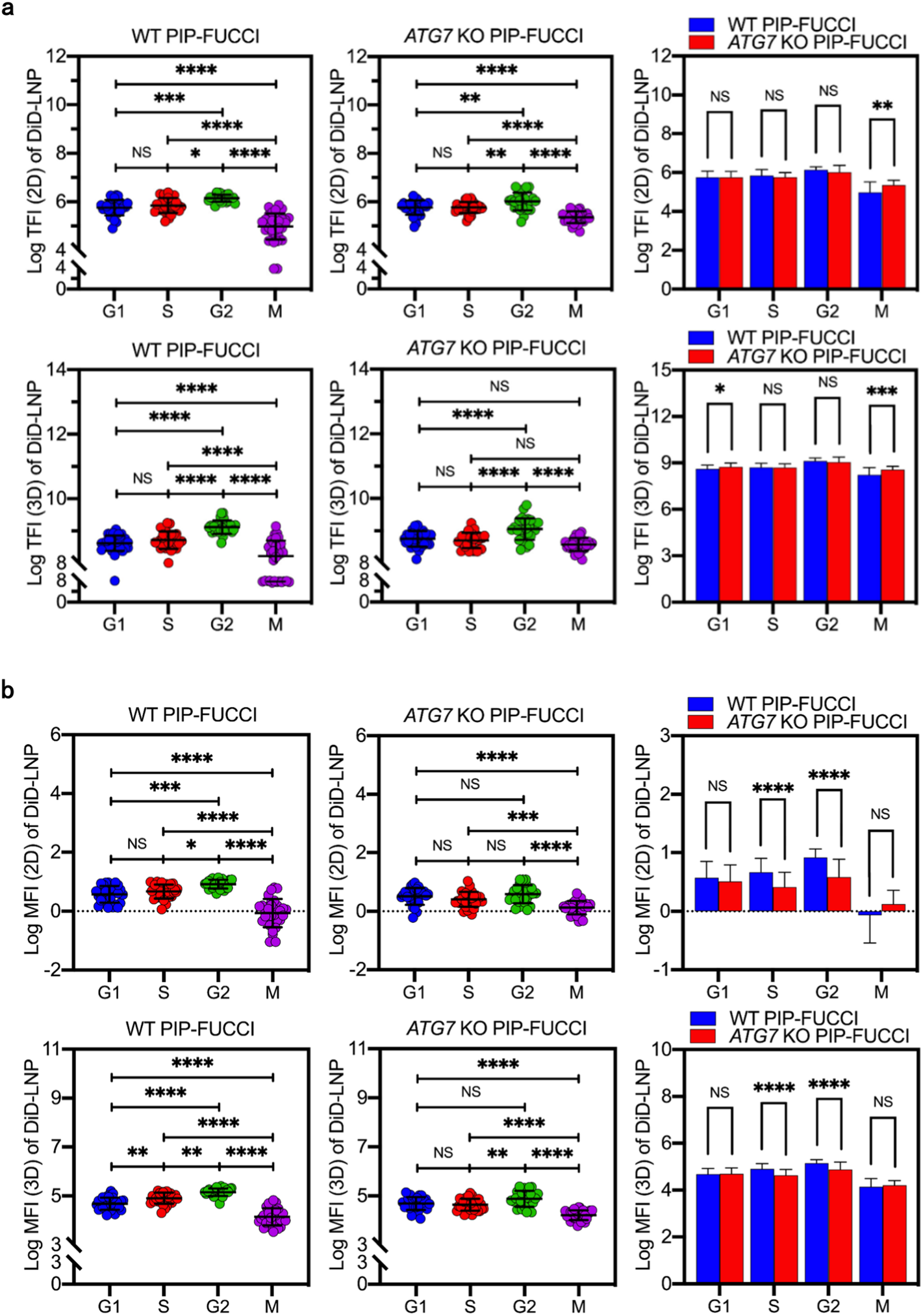
Determination of cell cycle- and endogenous autophagy-associated cellular accumulation of DiD-labelled lipid nanoparticles (DiD-LNP) using the newly established technical platform. **a**, Statistical analysis of Total Fluorescence Intensity (TFI) in G1-phase, S-phase, G2-phase and M-phase WT and *ATG7* KO PIP-FUCCI cells respectively measured at two-dimensional (2D, upper panel) and three-dimensional (3D, lower panel) levels. **b**, Statistical analysis of Mean Fluorescence Intensity (MFI) in WT and *ATG7* KO PIP-FUCCI cells respectively measured at four distinct cell cycle phases, as well as at 2D (upper panel) and 3D (lower panel) levels. WT PIP-FUCCI, wide type U2OS cells stably expressing PIP-FUCCI; *ATG7* KO PIP-FUCCI, *ATG7* knockout U2OS cells stably expressing PIP-FUCCI. DiD-LNP was administered at a dose equivalent to 6 μg/mL luciferase mRNA for 2 hours. All data were analyzed after log_10_ transformation and presented as “mean ± standard deviation” (n = 25-30). Statistical analysis of DiD-LIP accumulation among four distinct cell cycle phases was conducted using ANOVA with Bonferroni correction, while differences between WT and *ATG7* KO PIP-FUCCI cell lines were evaluated with an independent sample *t*-test. *, *P* < 0.05; **, *P* < 0.01; ***, *P* < 0.001; ****, *P* < 0.0001. NS, not significant.

### Characterization of fluorescence intensity-based parameters utilized in determination of cellular accumulation of DiD-LIP and DiD-LNP

Considering that TFI (2D), TFI (3D), MFI (2D) and MFI (3D) were firstly used in combination for determination of nanoparticle accumulation, their correlations were assessed with Spearman or Pearson Correlation Coefficients (see “Methods”). As indicated in Table 2, when the correlations between two dimensional levels were respectively evaluated for TFI and MFI in WT and *ATG7* KO PIP-FUCCI cells, moderate to very high Correlation Coefficients were obtained for all the assessed groups. Nevertheless, moderate to high Correlation Coefficients (*r* = 0.689 ∼ 0.793, all *P* < 0.0001) were achieved for TFI and MFI of DiD-LIP, while high to very high correlations (*r* = 0.811 ∼ 0.932, all *P* < 0.0001) were detected for those of DiD-LNP. Interestingly, further correlation analysis between TFI and corresponding MFI at same dimensional level revealed low to high Correlation Coefficients (*r* = 0.495 ∼ 0.711, all *P* < 0.0001) for TFI and MFI of DiD-LIP, but high to very high Correlation Coefficients (*r* = 0.791 ∼ 0.918, all *P* < 0.0001) for those of DiD-LNP (*r* = 0.791, *P* < 0.0001) (Table 3).

**Table 2.** Spearman Correlation Coefficient between TFI (2D) and TFI (3D), MFI (2D) and MFI (3D) in WT and *ATG7* KO PIP-FUCCI cell lines.

|  | WT PIP-FUCCI |  |  |  | *ATG7* KO PIP-FUCCI |  |  |  |
| --- | --- | --- | --- | --- | --- | --- | --- | --- |
|  | TFI (3D)<br>of DiD-LIP | TFI (3D)<br>of DiD-LNP | MFI (3D)<br>of DiD-LIP | MFI (3D)<br>of DiD-LNP | TFI (3D)<br>of DiD-LIP | TFI (3D)<br>of DiD-LNP | MFI (3D)<br>of DiD-LIP | MFI (3D)<br>of DiD-LNP |
| TFI (2D) of DiD-LIP | 0.791<br>($P < 0.0001$) | / | / | / | 0.715<br>($P < 0.0001$) | / | / | / |
| MFI (2D) of DiD-LIP | / | / | 0.793<br>($P < 0.0001$) | / | / | / | 0.689<br>($P < 0.0001$) | / |
| TFI (2D) of DiD-LNP | / | 0.872<br>($P < 0.0001$) | / | / | / | 0.811<br>($P < 0.0001$) | / | / |
| MFI (2D) of DiD-LNP | / | / | / | 0.932<br>($P < 0.0001$) | / | / | / | 0.905<br>($P < 0.0001$) |

**503** TFI, Total fluorescence intensity; MFI, Mean fluorescence intensity; DiD-LIP, DiD-labelled liposomes (at a dose equivalent to 5 µg/mL DOX); DiD-LNP, DiD-labelled lipid **504** nanoparticles (at a dose equivalent to 6 µg/mL luciferase mRNA); 2D, Two-dimensional level; 3D, Three-dimensional level. The level of significance was set at $P < 0.05$ .

**Table 3.** Spearman Correlation Coefficient between TFI (2D) and MFI (2D), TFI (3D) and MFI (3D) in WT and *ATG7* KO PIP-FUCCI cell lines.

|  | WT PIP-FUCCI |  |  |  | *ATG7* KO PIP-FUCCI |  |  |  |
| --- | --- | --- | --- | --- | --- | --- | --- | --- |
|  | MFI (2D)<br>of DiD-LIP | MFI (2D)<br>of DiD-LNP | TFI (3D)<br>of DiD-LIP | TFI (3D)<br>of DiD-LNP | MFI (2D)<br>of DiD-LIP | MFI (2D)<br>of DiD-LNP | TFI (3D)<br>of DiD-LIP | TFI (3D)<br>of DiD-LNP |
| TFI (2D) of DiD-LIP | 0.711<br>($P < 0.0001$) | / | / | / | 0.699<br>($P < 0.0001$) | / | / | / |
| MFI (3D) of DiD-LIP | / | / | 0.495<br>($P < 0.0001$) | / | / | / | 0.546<br>($P < 0.0001$) | / |
| TFI (2D) of DiD-LNP | / | 0.918<br>($P < 0.0001$) | / | / | / | 0.856<br>($P < 0.0001$) | / | / |
| MFI (3D) of DiD-LNP | / | / | / | 0.871<br>($P < 0.0001$) | / | / | / | 0.791<br>($P < 0.0001$) |

**505** TFI, Total fluorescence intensity; MFI, Mean fluorescence intensity; DiD-LIP, DiD-labelled liposomes (at a dose equivalent to 5 µg/mL DOX); DiD-LNP, DiD-labelled lipid **506** nanoparticles (at a dose equivalent to 6 µg/mL luciferase mRNA); 2D, Two-dimensional level; 3D, Three-dimensional level. The level of significance was set at $P < 0.05$ .

Importantly, further Statistical analysis using z-test on Fisher z-transformed correlation coefficients discovered that there were statistical significances between different nanoparticles and different cell lines on aforementioned Correlation Coefficients among TFI and MFI. For instance, Correlation Coefficients among TFI and MFI in DiD-LIP-treated groups were significantly lower than those treated with DiD-LNP in WT and *ATG7* KO PIP-FUCCI cells (*P* values, 0.0441 ∼ < 0.0001), except no correlation detected between TFI (2D) and TFI (3D) of DiD-LIP with that of DiD-LNP in *ATG7* KO PIP-FUCCI cells (*P* = 0.0860) (Supplementary Table 10). In addition, when comparing Correlation Coefficients among TFI and MFI in different cell lines, Correlation Coefficient between MFI (2D) and TFI (2D) of DiD-LNP in WT PIP-FUCCI cells was significantly higher than that in *ATG7* KO PIP-FUCCI cells (*P* = 0.0275) (Supplementary Table 11). These data suggest that nanoparticle type and *ATG7* expression status, particularly the former, may affect the correlation strength among fluorescence intensity-based parameters for nanoparticle accumulation determination.

On the other hand, as cell volumes gradually “grow” while cell cycle progressing, we further evaluated whether the increased fluorescence intensities were associated with increased cell volumes rather than cell cycle phases. Surprisingly, using TFI (3D) of DiD-LIP and DiD-LNP in WT and *ATG7* PIP-FUCCI cells as representative models, remarkable variations on Pearson Correlation Coefficients between TFI (3D) and cell volume were obtained among distinct cell cycle phases, as well as between different nanoparticles and cell lines (Supplementary Fig. 11 and Fig. 12). For instance, no correlation between TFI (3D) and cell volume was detected in the following groups: DiD-LIP-treated M-phase *ATG7* KO PIP-FUCCI cells, DiD-LNP-treated G1-phase WT PIP-FUCCI cells, as well as DiD-LNP-treated G1-phase, S-phase and G2-phase *ATG7* KO PIP-FUCCI cells, all *P* > 0.05. However, high Correlation Coefficients between TFI (3D) of DiD-LIP and cell volumes were achieved in WT PIP-FUCCI cells at of G1-phase (*r* = 0.808), G2-phase (*r* = 0.860) and M-phase (*r* = 0.805), while a low correlation coefficient (*r* = 0.493) was detected in S-phase WT PIP-FUCCI cells. Taken together, our data did not provide sufficient evidences on a clear correlation between fluorescence intensity-based parameters and cell sizes.

## Discussion

This study aims to evaluate the impact of cell cycle and endogenous autophagy on cellular accumulation of nanoparticles under conditions largely mimicking physiological status. Through construction and application of a platform simultaneously overcoming several long-lasting methodological obstacles, we present initial and multi**-**dimensional data demonstrating that G2-phase and M-phase cells respectively have the maximum and minimum accumulation of DiD-LIP and DiD-LNP (Fig. 4 and Fig. 5; Table 1). Nevertheless, these two distinct cell populations were previously muddled up together (so called G2/M-phase cells) because of methodological deficiencies in distinguishing one population from another without scathe. Moreover, differential accumulation of DiD-LIP and DiD-LNP was not only observed between spherical M-phase and adherent G2-phase cells, but also occurred among adherent cells in both WT and *ATG7* KO PIP-FUCCI cells. Interestingly, among adherent cells at three distinct cell cycle phases, mean values of TFI and MFI of both nanoparticles exhibited similar tendency in WT and *ATG7* KO PIP-FUCCI cells, which was G1 < S < G2. These findings may suggest a synchronization in intensified cellular metabolism and enhanced nanoparticle accumulation while cell cycle progressing from G1-phase to G2-phase. However, detailed mechanisms mediating cell cycle-associated nanoparticle accumulation warrant further explorations, particularly for “torpid” M-phase cells. On the other hand, our findings generally suggest an enhanced nanoparticle accumulation medicated by *ATG7* knockout, as demonstrated by TFI/MFI of LIP and TFI of LNP. Although further verification and exploration on underlying mechanisms are needed, our data may provide a novel line of supportive evidence for combined use of anticancer modalities with autophagy blockers currently under preclinical and clinical evaluations^52,53^.

It is noteworthy that in this cross-sectional study, cells representative for distinct cell cycle phases were randomly selected based on nuclear fluorescence and microscopic morphology. However, durations for each cell cycle phase have significant variations among different cell types and even within the same cell type cultured in different conditions. These variations are more pronounced in cancer cells due to their well-known heterogeneity. According to the known phase durations for mammalian cells and 2 hour-incubation timepoint designated, the representative cells selected in our study may include a majority of cells remaining in the same phase throughout the incubation period and a small portion progressed from the previous phase, particularly for G1-phase, S-phase and G2-phase cells. As this study intends to determine the intracellular nanoparticle levels for cells at four distinct cell cycle phases and with specific autophagy status respectively at the time of analysis, the net value added during each cell cycle phase has not been investigated herein. Nevertheless, acquiring time-lapse live cell imaging could help elucidating this interesting issue. Indeed, our preliminary longitudinal experiment conducted in WT and *ATG7* KO PIP-FUCCI cells monitored with CLSM for up to 25 hours has provided useful information and laid the foundation for further exploration on this topic (Supplementary Fig. 6).

A technical detail worth mentioning is that although the distinction between routinely growing adherent G2-phase and spherical M-phase cells based on morphology and attachment is straightforward in microscopy, in the present study, cultured cancer cells were firstly administered with fluorescence-labelled nanoparticles and EpCAM antibody. Afterwards, culture dishes underwent gentle rinses with PBS to remove unbounded nanoparticles and antibody. As suspended cells in culture medium will be directly removed with PBS rinses, mitotic shake-off was adopted to separately harvest mitotic cells. Subsequently, settled and concentrated mitotic cells were transferred to another culture dish after gentle rinses for image acquisitions, in parallel with adherent cells maintained in original culture dish (Fig. 2). In combination with cell cycle indicator PIP-FUCCI, this approach not only achieved thorough distinguishment of cells at four distinct cell cycle phases with minimal disturbance in physiological state, but enabled us to obtain a favorable number of mitotic cells. Meanwhile, increased efficiency and higher quality were achieved for time-sensitive image acquisition and 3D reconstruction with CLSM, as no interference existed among adherent and suspended cells.

To attenuate the potential influence of cell shape (particularly between cells at M-phase and other phases) and cell size, nanoparticle accumulation was determined at multi-dimensional levels, with TFI and MFI at both 2D and 3D levels were respectively calculated. Undoubtedly, integration of TFI (2D), TFI (3D), MFI (2D) and MFI (3D), four fluorescence intensity-based and correlated parameters as assessed with Spearman Correlation Coefficients will promote more comprehensive and precise understandings on cell cycle- and autophagy-associated nanoparticle accumulation (Table 2 and Table 3). Interestingly, if cells and nanoparticles were respectively considered as individuals and therapeutic drugs, cell size-related and fluorescence intensity-based parameters may remind us of similar parameters used in clinical practice alone or in combination, e.g. total body surface area, mean body surface area, total body weight, body mass index, as well as drug exposure- and drug dose-related parameters. On the other hand, as 3D assessment offers a more comprehensive consideration of various factors, e.g. morphological changes during cell cycle progression, disparities in cellular volume^45,46^ and non-uniform intracellular distribution of nanoparticles, 3D-based parameters might aid in mitigating the constraints associated with 2D level and facilitates the “correction” of experimental findings. Further assessment of these parameters based on a large amount of application data will not only deepen our understandings on their values and significances, but also promote optimization of the technical platform constructed in this study. Finally, although several issues warrant further investigation, e.g. detailed mechanisms mediating cell cycle- and autophagy-associated nanoparticle accumulation, possible interplay between cell cycle and autophagy, merits of TFI and MFI at different dimensional levels, as well as possible nanoparticle- and cell line-mediated specificities on nanoparticle accumulation, our findings may open new insights into optimizing the design and application of lipid-based and possibly other nanomedicines besides providing a valuable technical solution. Particularly, around 3,200 LIP/LNP-based nano-formulations are currently under clinical investigations. In addition, it is also notable that the technical platform constructed in this study may provide a new paradigm or a new strategy for simultaneous investigation on other “effectors” in addition to cell cycle, or on the basis of complete cell cycle phase separation.

## Supporting information

Supplementary Video 1 (3D DiD-LlP accumulation in G1-phase WT PIP-FUCCI cells).

Supplementary Video 2 (3D DiD-LlP accumulation in S-phase WT PIP-FUCCI cells).

Supplementary Video 3 (3D DiD-LlP accumulation in G2-phase WT PIP-FUCCI cells).

Supplementary Video 4 (3D DiD-LlP accumulation in M-phase WT PIP-FUCCI cells).

Supplementary Video 5 (3D DiD-LlP accumulation in G1-phase ATG7 KO PIP-FUCCI cells)

Supplementary Video 6(3D DiD-LlP accumulation in S-phase ATG7 KO PIP-FUCCI cells).

Supplementary Video 7 (3D DiD-LlP accumulation in G2-phase ATG7 KO PIP-FUCCI cells).

Supplementary Video 8 (3D DiD-LlP accumulation in M-phase ATG7 KO PIP-FUCCI cells).

Supplementary Video 9 (3D DiD-LNP accumulation in G1-phase WT PIP-FUCCI cells).

Supplementary Video 10 (3D DiD-LNP accumulation in S-phase WT PIP-FUCCI cells).

Supplementary Video 11 (3D DiD-LNP accumulation in G2-phase WT PIP-FUCCI cells).

Supplementary Video 12 (3D DiD-LNP accumulation in M-phase WT PIP-FUCCI cells).

Supplementary Video 13 (3D DiD-LNP accumulation in G1-phase ATG7 KO PIP-FUCCI cells).

Supplementary Video 14 (3D DiD-LNP accumulation in S-phase ATG7 KO PIP-FUCCI cells).

Supplementary Video 15 (3D DiD-LNP accumulation in G2-phase ATG7 KO PIP-FUCCI cells).

Supplementary Video 16 (3D DiD-LNP accumulation in M-phase ATG7 KO PIP-FUCCI cells).

## Methods

### Major reagents

Cholesterol, 1-palmitoyl-2-stearoyl-sn-glycero-3-phosphocholine (HSPC), 1,2-distearoyl-sn-glycero-3-phosphocholine (DSPC) and 1,2-distearoyl-sn-glycero-3-phosphoethanolamine-N-[methoxy(polyethylene glycol)-2000] (mPEG_2000_-DSPE) were from Lipoid GmbH (Ludwigshafen, Germany). [(4-Hydroxybutyl)azanediyl]di(hexane-6,1-diyl) bis(2-hexyldecanoate) (ALC-0315) and 2-[(polyethylene glycol)-2000]-N,N-ditetradecylacetamide (ALC-0159) were acquired from SINOPEG (Xiamen, China). 1,1′-Dioctadecyl-3,3,3′,3′-tetramethylindotricarbocyanine, 4-chlorobenzenesulfonate salt (DiD) was from MKBio (Shanghai, China). Firefly luciferase mRNA (LUC mRNA) was from Afana Biotechnology (Hefei, China). Ferric chloride hexahydrate and ammonium thiocyanate were from Sigma-Aldrich (St. Louis, MO, USA). Lipofectamine 3000 and Opti-MEM were purchased from Thermo Fisher Scientific (Waltham, MA, USA). Polybrene was from Santa Cruz (Dallas, TX, USA). G418 was from InvivoGen (San Diego, CA, USA). Glutaraldehyde, ethanol and acetone were from Hushi Chemical (Shanghai, China). Lead citrate was from Macklin (Shanghai, China). RIPA buffer and DAPI staining solution were from Beyotime (Shanghai, China). Proteases inhibitor and phosphatase inhibitor were from Yeason (Shanghai, China). Protein marker was from BioRad (Hercules, CA, USA).

Lentivirus concentration kit (Cat. NO. 41101ES50) was acquired from Yeason. BCA protein assay kit (Cat. NO. FD2001) was obtained from Thermo Fisher Scientific. E.Z.N.A. Endo-free plasmid maxi kit (Cat. NO. D6926-03) was from Omega (Norcross, GA, USA). Plasmids of psPAX2 (Cat. NO. 12260), pMD2.G (Cat. NO. 12259), pLenti-PGK-Neo-PIP-FUCCI (PIP-FUCCI) (Cat. NO. 118616) and pLenti PGK Neo DEST (empty vector) (Cat. NO. 19067) were from Addgene (Cambridge, MA, USA).

Anti-ATG7 antibody (Cat. NO. ab133528; clone name, EPR6251) and anti-SQSTM1/p62 antibody (Cat. NO. ab91526) were from Abcam (Waltham, MA, USA). LC3A/B XP Rabbit mAb (Cat. NO. 12741S; clone name, D3U4C) and β-actin antibody (Cat. NO. 4967) were from Cell Signaling Technology (Danvers, MA, USA). EpCAM monoclonal antibody, FITC (Cat. NO. MA1-10197; clone name, VU-1D9), goat anti-rabbit IgG (H+L) highly cross-adsorbed secondary antibody (Alexa Fluor 488) (Cat. NO. A32731), goat anti-rabbit IgG (H+L) cross-adsorbed secondary antibody (Alexa Fluor 546) (Cat. NO. A11010), and goat anti-mouse IgG (H+L) cross-adsorbed secondary antibody (Alexa Fluor 546) (Cat. NO. A11003) were from Thermo Fisher Scientific.

### Cell lines and cell cultures

Wild type (WT) and *ATG7* knockout (*ATG7* KO) human osteosarcoma cancer cell lines U2OS were kind gifts from Professor Qiming Sun at Zhejiang University School of Medicine. Human kidney epithelial cell line 293T was from the Shanghai Institute of Cells, Chinese Academy of Sciences. All cell lines, including newly established WT and *ATG7* KO PIP-FUCCI cell lines (see below section), were authenticated by STR profiling (Tsingke Biotechnology, Beijing, China) and confirmed to be mycoplasma free. All cell lines were in cultured in DMEM supplemented with 10% (v/v) fetal bovine serum and maintained at 37 °C in an atmosphere containing 5% CO_2_.

### Preparation of model nanoparticles DiD-LIP and DiD-LNP

DiD-LIP was synthesized according to the officially published formulation of Doxil^®35^ and previous reports^54^. First, HSPC, cholesterol and mPEG_2000_-DSPE, at 56.6:38.1:5.3 molar ratio, were hydrated in 10 mM sucrose-histidine buffer at pH6.5. Subsequently, DiD was added into the lipid mixture at a concentration of 0.4 mol%, followed by vortex at 60 °C for 2-3 minutes to generate multilamellar vesicles (MLVs). Finally, the MLVs were reduced in size using 400, 100, 80, and 50 nm-pore size polycarbonate filters with a 10-mL extruder barrel (Northern Lipids, Vancouver, Canada), and stored at a temperature of 2-8 °C until use.

DiD-LNP was prepared based on the officially published formulation of Comirnaty^®36^ using a modified procedure we previously described^4,55^. Initially, ALC-0315, DSPC, cholesterol and ALC-0159 were dissolved at a molar ratio of 46.3: 9.4: 42.7: 1.6 to fabricate the ethanol phase and DiD was then added into the lipids at a concentration of 0.4 mol%. Meanwhile, LUC mRNA was dissolved in 20 mM citrate buffer to prepare the aqueous phase. Subsequently, the above two phases were mixed with a microfluidic mixer (Precision Nanosystems, Canada) at a flow rate ratio of 1:3 (ethanol: aqueous). Afterwards, the resulting nanoparticle solution was subjected to dialysis against 10 × volume of PBS for at least 18 hours using tangential flow filtration (TFF) membranes with molecular weight cut-offs of 100 kD (Sartorius Stedim, Germany). With the assistance of an Amicon ultracentrifugal filter (EMD Millipore, Billerica, MA, USA), the nanoparticle solution was passed through a 0.22 µm filter and ultimately concentrated to the desired concentration.

### General characterization of DiD-LIP and DiD-LNP

As we previously introduced in details^4^, a Zetasizer Nano ZS instrument (Malvern Instruments Ltd, Malvern, UK) was employed to measure hydrodynamic size (Z-average), polydispersity index (PDI) and zeta potential of DiD-LIP and DiD-LNP, with the temperature maintained at 25 °C throughout the testing period. To further evaluate their stability in experimental environment, DiD-LIP and DiD-LNP were diluted with cell culture solution and incubated at 37 °C for 24 hours. Afterwards, nanoparticle samples were respectively collected at specified time points (1 hour, 2 hours, 6 hours, 12 hours and 24 hours post-incubation). Except that the test temperature was changed to 37 °C, samples collected were subjected to aforementioned characterization of Z-average and PDI. Moreover, morphology of DiD-LIP and DiD-LNP was examined with Cryo-TEM, with images captured with a Talos F200C instrument (FEI/Thermo Scientific, Waltham, MA, USA) equipped with a Ceta 4k × 4k camera at an acceleration voltage of 200 kV. Moreover, the phospholipid (HSPC and mPEG_2000_-DSPE) concentration of DiD-LIP solutions and phospholipid (DSPC) concentration of DiD-LNP solutions were determined through Steward’s assay as previously described in details^4,56,57^. Eventually, calibration curves were established for DSPC lipids within the range of 25 to 200 μg/mL, and for HSPC lipids within the range of 25 to 150 μg/mL. These curves were applied to determine the ultimate concentrations of phospholipids in the DiD-LIP and DiD-LNP solutions. Three separate experiments were conducted for further statistical analysis. According to previous pharmacokinetics studies and related literatures, the doses of DiD-LIP and DiD-LNP used in this study were equivalent to 5 μg/mL DOX ^35,58,59^ (42.67 μg/mL of lipids including 25.61 μg/mL of HSPC, 8.53 μg/mL of mPEG_2000_-DSPE and 8.53 μg/mL of Cholesterol) and 6 μg/mL LUC mRNA ^60–62^ (154.20 μg/mL of lipids including 85.80 μg/mL of ALC-0315, 10.20 μg/mL of ALC-0159, 18.00 μg/mL of DSPC and 40.20 μg/mL of Cholesterol), respectively. Same volume of PBS was used as a control.

### Construction of isogenic WT and *ATG7* KO PIP-FUCCI cell lines stably expressing cell cycle indicator PIP-FUCCI

As previously reported, in the cell cycle indicator plasmid PIP-FUCCI, mVenus fluorescent protein was fused to the PCNA-interacting protein (PIP) degron of human *Cdt1* gene (*Cdt1*_1-17_), which is rapidly degraded at the onset of DNA replication^33^. Meanwhile, mCherry fluorescent protein was fused to the *Geminin* gene (*Gem*_1-110_), which accumulates during S-phase, G2-phase and M-phase and begins to be degraded at late M-phase^33,63^. Based on these characteristics, Grant et al. demonstrated that in U2OS cell line transfected with PIP-FUCCI, mVenus single-positive cells displaying green nuclear fluorescence are at G1-phase, mCherry single-positive cells displaying red nuclear fluorescence are at S-phase, while mVenus and mCherry double-positive cells displaying yellow (overlap of green and red) nuclear fluorescence are at either G2-phase or M-phase^33^.

To take the advantage of cell cycle indicating capability of PIP-FUCCI plasmid in living cells, hereby WT and *ATG7* KO U2OS cell lines were stably transfected with PIP-FUCCI. Briefly, 1 × 10^6^ 293T cells were cultured into a 10-cm dish for overnight incubation. For lentiviral transduction, Lipofectamine 3000 was used to co-transfect PIP-FUCCI plasmid, psPAX2 and pMD2.G plasmids into 293T cells, with empty vector serving as a negative control. The medium was replaced 6-8 hours after transfection, and was harvested and concentrated at 72 hours post-transfection. Afterwards, approximately 6 × 10^4^ exponentially growing WT and *ATG7* KO U2OS cells were respectively seeded into cell culture plates. After overnight incubation, cells were exposed to concentrated lentiviral supernatant containing either PIP-FUCCI or empty vector, along with 8 μg/mL polybrene. Transduced cells were then selected for around one week with 500 μg/mL of G418. WT and *ATG7* KO U2OS cells expressing the empty vector, so called WT and *ATG7* KO Vector cell lines respectively, were amplified for subsequent experiments, while those transfected with PIP-FUCCI were sorted by BD FACSAria II flow cytometry (BD Biosciences, CA, USA) to obtain WT and *ATG7* KO PIP-FUCCI cell lines exhibiting strong mVenus and/or mCherry fluorescence for further studies.

### Cell-cycle phase tracking of WT and *ATG7* KO PIP-FUCCI cells through time-lapse live-cell imaging

Approximately 8 × 10^4^ exponentially growing WT and *ATG7* KO PIP-FUCCI cells were seeded into confocal dishes for overnight incubation. Live-cell imaging was conducted using an Andor BC43 spinning disk confocal microscope (Oxford Instruments, Abingdon, UK) at 20 × magnification (0.8 NA) for a duration of up to 25 hours, corresponding to the estimated cell doubling time (Supplementary Fig. 5). The imaging was performed under controlled conditions of 5% CO_2_ and 37 °C, with images acquired at 5-minute intervals.

### Evaluation of the influence of PIP-FUCCI transfection on cell morphology and proliferation

WT and *ATG7* KO Vector cell lines, as well as WT and *ATG7* KO PIP-FUCCI cell lines were carefully checked with light microscope for morphological examination. In addition, these cell lines were respectively seeded into 6-well plates (1 × 10^5^ cells for each well). A Countess II FL Automated Cell Counter (Thermo Fisher Scientific) was employed for five consecutive cell number counting between 24 hours and 120 hours after seeding, with a 24-hour interval. Cell doubling time of each cell line was calculated according to the following equation: Doubling time = Duration time of cell culture × log_10_ (2)/[log_10_ (final cell number) – log_10_ (initial cell number)].

### Cell cycle distribution detected by flow cytometry

Exponentially growing WT and *ATG7* KO PIP-FUCCI cells were seeded at a density of 1 × 10^6^ cells per 10-cm dish and cultured overnight. Cells were then respectively administered with DiD-LIP and DiD-LNP for 2 hours. Afterwards, cells were first gently shaken (so called mitotic shake-off) and the culture medium containing floating mitotic cells was harvested separately. Then, the remaining adherent G1-phase, S-phase and G2-phase cells were harvested after digestion with trypsinization. In addition, exponentially growing WT and *ATG7* KO PIP-FUCCI cells were seeded at 3 × 10^5^ cells per well into 6-well plates and cultured overnight. Cells were then respectively administered with DiD-LIP and DiD-LNP for 12 hours, 24 hours and 48 hours. Afterwards, cells were routinely harvested with trypsinization, rinsed three times with PBS. Each cell sample was suspended in 1 mL of DAPI staining solution and subsequently analyzed in equal volumes with a CytoFLEX LX (Beckman Coulter, Indianapolis, USA) to determine the cell cycle distribution. DAPI, mVenus and mCherry channels were respectively excited with 375 nm, 488 nm and 561 nm lasers, and detected with corresponding 450/45 nm, 525/40 nm and 610/20 nm bandpass filters. As described above, cancer cells exhibited the following cell cycle-dependent fluorescence: G1-phase (mVenus positive), green fluorescence; S-phase (mCherry positive), red fluorescence; G2-phase and M-phase (mVenus and mCherry double-positive), yellow fluorescence.

### Examination of possible effects of nanoparticles on autophagic vesicle formation with TEM

Exponentially growing WT and *ATG7* KO PIP-FUCCI cells were seeded at 7 × 10^5^ cells per 6-cm culture dish for overnight incubation. Cells were then respectively administered with DiD-LIP and DiD-LNP for 2 hours. Following overnight fixation with 2.5% glutaraldehyde, cells underwent three washes with PBS for a duration of 10 minutes each. Subsequently, cells were subjected to fixation with 1% osmium tetroxide for 1 hour, followed by rinsing with PBS prior to staining with 2% uranyl acetate for 30 minutes. For dehydration, samples were respectively placed in 50%, 70%, 90% and 100% ethanol for 15 minutes and then in 100% acetone for 20 minutes twice. Afterwards, a Leica UC7 microtome (Leica, Solms, Germany) was used to prepare ultrathin sections and sample staining were performed with lead citrate and uranyl acetate. Photographs of autophagic vesicles were captured using a Tecnai 10 TEM (Philips, Amsterdam, Netherlands) at 100 kV. At least 10 cells were randomly chosen from each group for imaging and statistical analysis.

### Western blotting to assess possible effects of nanoparticles on autophagy-related proteins

Exponentially growing WT and *ATG7* KO PIP-FUCCI cells were seeded at 7 × 10^5^ cells per 6-cm culture dish for overnight incubation. Cells were then respectively administered with DiD-LIP and DiD-LNP for 2 hours. Then cells were harvested with trypsinization, rinsed three times with PBS and subjected to RIPA buffer containing proteases and phosphatase inhibitors on ice for 10 minutes. Subsequently, the supernatant was collected through centrifugation at 12,000 × g for 20 minutes. Protein concentrations were determined with a BCA protein assay kit. A gel of 15% SDS-PAGE was used to electrophorese proteins (40 μg /lane) prior to being transferred onto a PVDF membrane. After blocking with 5% (w/v) bovine serum albumin for 1 hour, the membranes were incubated with following antibodies overnight at 4 °C: anti-ATG7 (1:1,000 dilution), anti-SQSTM1/p62 (1:1,000 dilution), anti-MAP1LC3A/B (1:1,000 dilution) and anti-β-actin (1:5,000 dilution). Next, membranes were rinsed with TBS solution containing 0.1% (v/v) Tween 20 and incubated with corresponding secondary antibodies (1:5,000 dilution) for 2 hours. Odyssey imaging system was employed for WB imaging (LI-COR Biosciences, Lincoln, USA).

### Integration of PIP-FUCCI expression with mitotic shake-off for distinguishment and collection of cells at four cell cycle phases and determination of cell cycle-associated nanoparticle accumulation

Approximately 8 × 10^4^ exponentially growing WT and *ATG7* KO PIP-FUCCI cells were seeded into confocal dishes for overnight incubation. Then cells were administered with DiD-labelled model nanoparticles (blue intracellular fluorescence in CLSM) at designated doses and incubated for 1.5 hours. Afterwards, cells were incubated with 25 μL FITC-EpCAM monoclonal antibody for additional 30 minutes to draw the outline of cell membrane (green cell membrane fluorescence in CLSM) and meanwhile exclude apoptotic cells or cell debris.

It should be noted that the use of PBS rinses to remove above unbound nanoparticles and antibody prior to subsequent nanoparticle fluorescence detection results in the loss of a large proportion of M-phase cells. Fortunately, unique properties of typical M-phase cells, e.g. spherical shape and significantly decreased adhesion capability^41^ facilitate convenient separation by gentle mitotic shake-off^34^, through which a complete distinguishment and adequate collection of cells at four cell cycle phases were successfully achieved by integrating PIP-FUCCI with mitotic shake-off in this study.

To achieve complete separation and adequate collection of cells at four different cell cycle phases, cells cultured in confocal dishes were gently shaken, and the culture medium containing floating mitotic cells was harvested with sterile centrifuge tubes, then settled and concentrated mitotic cells were transferred into another new confocal dish. Both the adherent G1-phase (mVenus single-positive, green nuclear fluorescence in CLSM), S-phase (mCherry single-positive, red nuclear fluorescence in CLSM) and G2-pahse (mVenus and mCherry double-positive, yellow nuclear fluorescence in CLSM) cells remained in original confocal dishes, and the floating M-phase cells (mVenus and mCherry double-positive, yellow nuclear fluorescence in CLSM) harvested separately via mitotic shake-off into new dishes were gently rinsed three times with PBS to remove unbound fluoroprobes, before examination with high-resolution CLSM and 3D reconstruction technique (see below section). It is noteworthy that although the emission wavelength of FITC partially overlaps with that of mVenus, FITC-labeled EpCAM expresses on the cell membrane as an epithelial cell adhesion molecule^64^, whereas mVenus expresses in the nucleus^33^. Consequently, there is no interference between them due to distinct expression locations.

### High-resolution CLSM and 3D reconstruction technique for multi-dimensional evaluation of cell cycle- and autophagy-associated nanoparticle accumulation

Successful construction of WT and *ATG7* KO PIP-FUCCI cell lines and achievement of complete cell cycle phase separation have provided us with the opportunity to simultaneously determine cell cycle- and autophagy-associated cellular accumulation of DiD-labelled model nanoparticles. Live-cell imaging was conducted using the Confocal Zeiss LSM 880 (Carl Zeiss, Jena, Germany) equipped with a 63 × oil immersion objective (1.40 NA) with a resolution of 2048 × 2048 pixels, together with an incubation chamber set at 37 °C and 5% CO_2_. Filter sets were as follows: mVenus (excitation: 514 nm, emission: 519–590 nm), mCherry (excitation: 561 nm, emission: 570–620 nm), DiD (excitation: 633 nm, emission: 638–756 nm) and FITC (excitation: 488 nm, emission: 500–550 nm)^33^. In addition to labelling cell membrane with FITC, morphological examination of cell with bright field was helpful in identifying typical and healthy M-phase cells with intact and smooth cell membranes.

Besides regular two-dimensional (2D) CLSM imaging to determine the cell cycle phase and nanoparticle accumulation, three-dimensional (3D) imaging was also conducted by acquiring 30-80 μm z-stacks at z-step intervals of 0.4-1 μm. DiD-LIP and DiD-LNP were respectively added into cells, using same volume of PBS as a control. DiD fluorescence was detected under identical conditions for both control groups and nanoparticle-treated groups, with background intensity determined according to DiD fluorescence detected in corresponding control groups. Following delineation of cell boundaries in 2D images using brightfield and FITC fluorescence, TFI of intracellular DiD was quantified using Fiji/Image J (imagej.nih.gov). Subtraction of the background intensity yielded TFI (2D) of DiD in experimental cells. To eliminate variations in TFI (2D) due to differences in cell area, mean fluorescence intensity (MFI) of intracellular DiD at the 2D level, so called MFI (2D), was calculated as the ratio of TFI (2D) vs. cell area^65^. Imaris software (Bitplane) was used to reconstruct 3D images of confocal z-stacks, with a series of parameters such as Surface Grain Size and Manual Threshold Value parameters maintained consistent during signal processing to determine TFI of intracellular DiD. Similarly, TFI (3D) was obtained based on TFI of DiD subtracting the background intensity To eliminate variations in TFI (3D) due to differences in cell volume, TFI (3D) was divided by corresponding cell volume to obtain MFI (3D)^66^.

### Correlation analysis and comparison of correlations among groups

Pearson correlation coefficients were employed to evaluate the correlations between TFI (3D) and cell volume, and Spearman correlation coefficients were employed to evaluate the correlations between multi-dimensional TFI and MFI parameters. Correlations with *P* < 0.05 were considered significant and the degree of correlation was assessed using correlation coefficient (*r*) (*r* values, 0.00 ∼ < 0.03, negligible correlation; 0.30 ∼ < 0.50, low correlation; 0.50 ∼ < 0.70, moderate correlation; 0.70 ∼ < 0.90, high correlation and 0.90 ∼ < 1.00, very high correlation)^67^. In addition, comparison of correlations among groups was compared with a z-test on Fisher z-transformation, and *P* < 0.05 was considered significant^68^.

### Data presentation and statistical analysis

Data were presented as “mean ± standard deviation”. Since the normality test for continuous parametric data is a prerequisite for determining the statistical analysis method^4,69,70^, Shapiro-Wilk test was used to evaluate normality. Nonparametric tests generally have lower statistical power compared to parametric tests, especially in studies with small sample sizes^71^. Therefore, when the normal distribution of original or transformed data were confirmed, parametric tests were conducted. Otherwise, nonparametric tests were used. Parametric tests, One-way analysis of variance (ANOVA) with Bonferroni correction and an independent *t*-test were conducted for analyzing possible changes on nanoparticle stability and cell cycle distribution. Mann-Whitney U test (nonparametric test) was employed to analyze statistical differences in autophagic vesicle number among various groups. In addition, the raw data of cell area, cell volume, TFI and MFI of DiD-LIP/DiD-LNP at different dimensional levels, exhibited a partially non-normal distribution. However, they became a normal distribution after log_10_ transformation. Thus, the above raw data were transformed prior to statistical analyses, their differences among four distinct cell cycle phases were conducted using ANOVA with Bonferroni correction, and their differences between WT and *ATG7* KO PIP-FUCCI cell lines were evaluated with an independent sample *t*-test. All statistical analyses were conducted using SPSS 21.0 (IBM SPSS, Chicago, USA). *P* < 0.05 is considered statistically significant.

## Acknowledgments

This work was supported by grants from Zhejiang Provincial Natural Science Foundation of China (LZ23E030003 to Dr. Meihua Sui), the National Natural Science Foundation of China (21722405 and 22075243 to Dr. Meihua Sui) and Startup Foundation for Hundred-Talent Program of Zhejiang University (to Dr. Meihua Sui). We would like to acknowledge Dr. Yingjie Wang, Dr. Wei Liu, Dr. Yongzhong Du and Dr. Qiming Sun at Zhejiang University for insightful discussions. In addition, we would like to acknowledge Shenghai Chang, Lingyun Wu and Beibei Wang in the Center of Cryo-Electron Microscopy (CCEM), Zhejiang University for their technical assistance on Cryo-EM and TEM. We would like to acknowledge Shuangshuang Liu, Guifeng Xiao and Yueting Xing from the Core Facilities, Zhejiang University School of Medicine for their technical assistance on confocal live imaging, 3D reconstruction and flow cytometry. We would also like to acknowledge Yina Wang for her assistance on language polishing.

## Author contributions

M.S., the principal investigator of the major supporting grants, and Y.W. conceived the study and defined the goals of the present study. M.S., Y.W., G.L. and H.W. designed the experiments. Y.W., G.L. and H.W. performed the research. X.X. provided technical/platform supports. M.S., Y.W., G.L., H.W., Y.Z., W.Z., J.L., B.C. and Y.J. discussed this project and analyzed the data. M.S., Y.W., G.L. and H.W. drafted the manuscript. M.S., Y.W., G.L., H.W., Y.Z., W.Z., J.L., B.C. and Y.J. revised and edited the manuscript. These authors contributed equally: Y.W., G.L. and H.W.

## Data availability

All data supporting the findings of this study are available within the paper and Supplementary Information. The associated raw data are available from the corresponding author on reasonable request.

**Correspondence and requests for materials** should be addressed to Meihua Sui.

