## Supplementary materials for "Construction and application of a technical platform for determining cell cycle- and autophagy-associated cellular accumulation of lipid-based nanoparticles"

### Supplementary Information

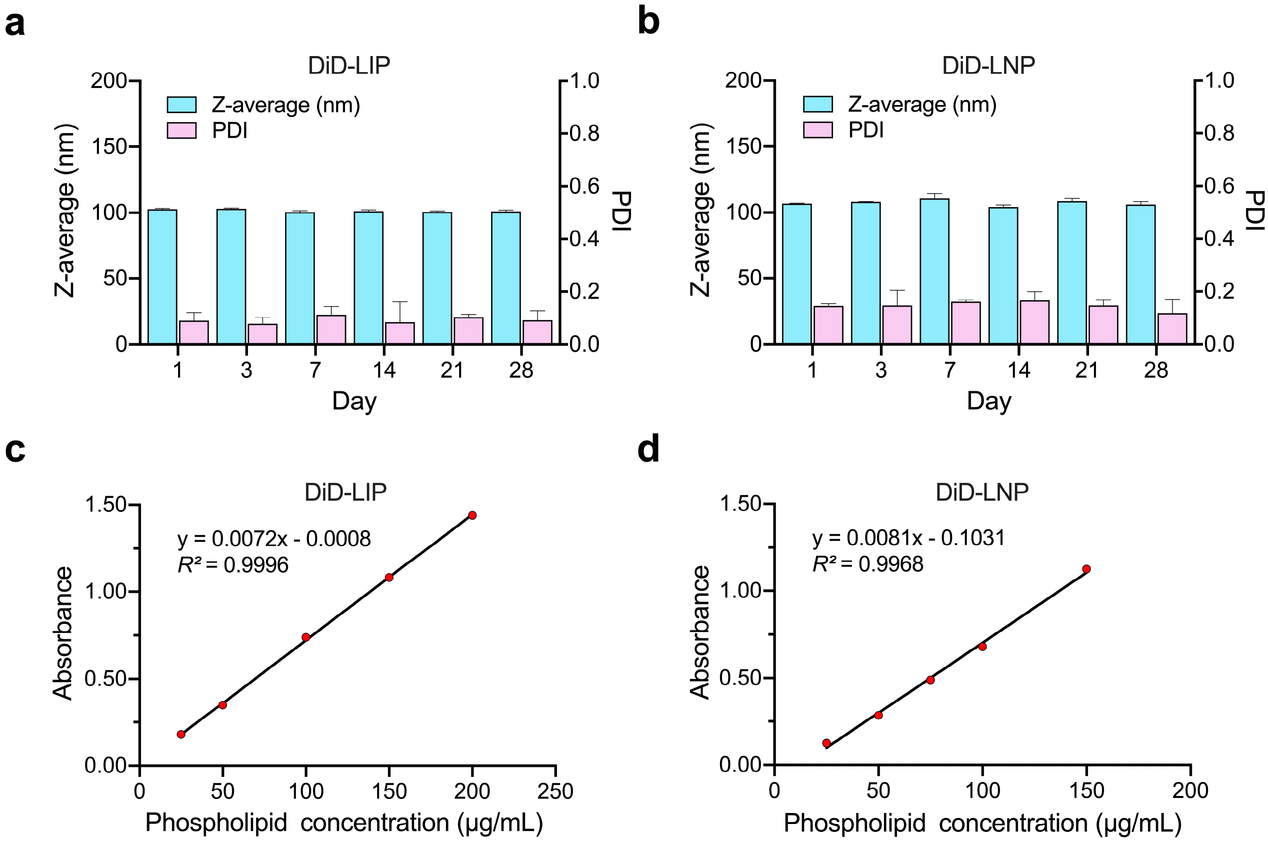

**Supplementary Fig. 1**. **Stability stored at 4 °C of DiD-labelled liposomes (DiD-LIP) and lipid nanoparticles (DiD-LNP), and standard curves for phospholipid. a-b,** Stability stored at 4 °C. DiD-LIP and DiD-LNP were respectively diluted to 1:100 with PBS and stored at 4°C for up to 28 days. Subsequently, 1 mL of nanoparticle solution was collected at designated time points (1 day, 3 days, 7 days, 14 days, 21 days and 28 days), followed by characterization of Z-average and PDI with dynamic light scattering. Z-average/PDI at six successive time points were as follows: DiD-LIP, 102.667 ± 0.737 nm/0.090 ± 0.031, 102.967 ± 0.666 nm/0.079 ± 0.024, 100.500 ± 0.964 nm/0.112 ± 0.031, 101.167 ± 0.987 nm/0.085 ± 0.077, 100.733 ± 0.569 nm/0.104 ± 0.010 and 101.000 ± 0.954 nm/0.092 ± 0.036; DiD-LNP, 106.767 ± 0.306 nm/0.145 ± 0.009, 108.167 ± 0.153 nm/0.147 ± 0.057, 110.700 ± 3.897 nm/0.161 ± 0.007, 104.300 ± 1.493 nm/0.166 ± 0.033, 108.700 ± 1.929 nm/0.147 ± 0.022 and 106.100 ± 2.265 nm/0.118 ± 0.052. Data were presented as "mean ± standard deviation" of three independent experiments. Z-average and PDI among various time points were evaluated by ANOVA with Bonferroni correction, with no significant difference detected (all $P > 0.05$). c-d, Standard curves for determining phospholipid (HSPC and $mPEG_{2000}$-DSPE in DiD-LIP, DSPC in DiD-LNP) concentration. The following equations were respectively obtained, in which y represented the absorbance measured at 470 nm and x represented the phospholipid concentration: DiD-LIP, $y = 0.0072x - 0.0008$ ($R^2 = 0.9996$); DiD-LNP, $y = 0.0081x - 0.1031$ ($R^2 = 0.9968$).

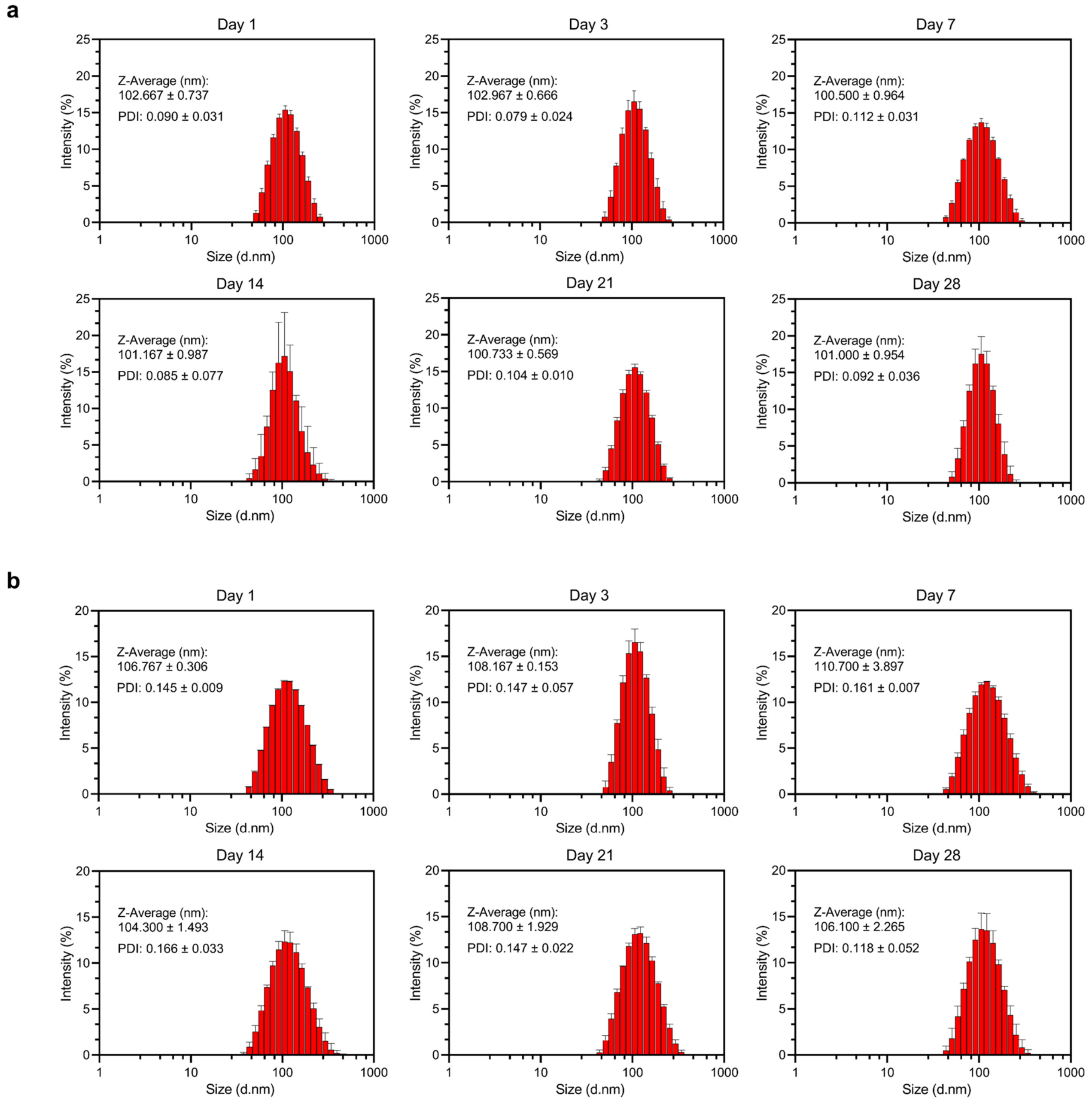

**Supplementary Fig. 2. Stability of DiD-labelled liposomes (DiD-LIP) and lipid nanoparticles (DiD-LNP) in PBS. a-b,** DiD-LIP (a) and DiD-LNP (b) were respectively diluted to 1:100 with PBS and stored at 4 ℃ for up to 28 days. Subsequently, 1 mL of nanoparticle solution was collected at designated time points (1 day, 3 days, 7 days, 14 days, 21 days and 28 days), followed by characterization of Z-average and PDI with dynamic light scattering. Data were presented as "mean ± standard deviation" of three independent experiments.

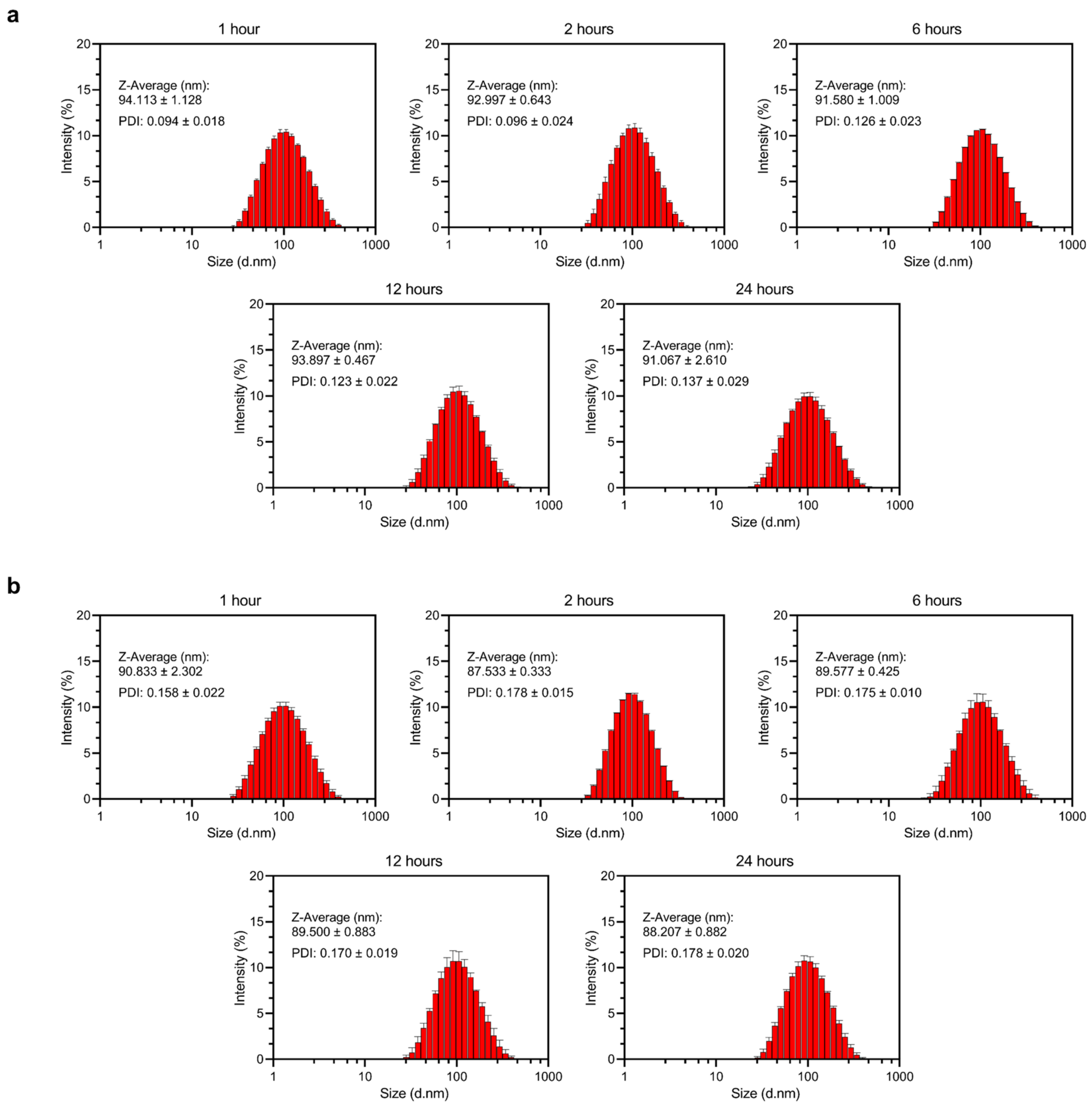

**Supplementary Fig. 3. Stability of DiD-labelled liposomes (DiD-LIP) and lipid nanoparticles (DiD-LNP) in serum. a-b,** DiD-LIP (a) and DiD-LNP(b) were respectively diluted to 1:100 with DMEM containing 10% FBS and incubated at 37 °C for 24 hours. Subsequently, 1 mL of diluted nanoparticle solution was collected at designated time points (1 hour, 2 hours, 6 hours, 12 hours and 24 hours), followed by characterization of Z-average and PDI with dynamic light scattering. Data were presented as "mean ± standard deviation" of three independent experiments.

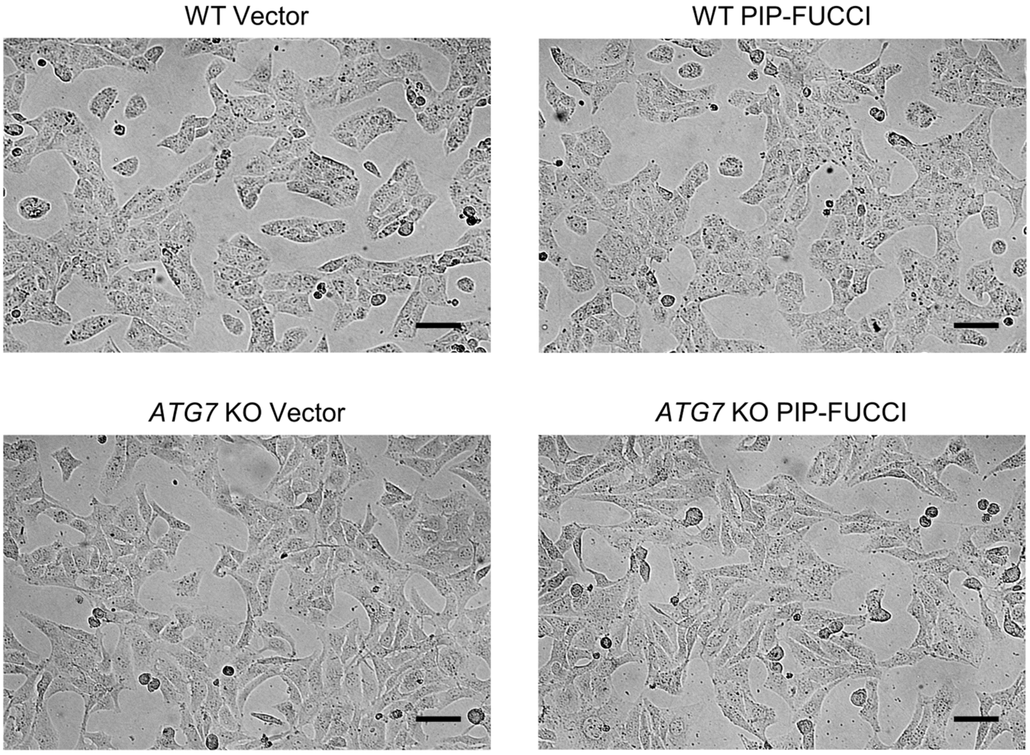

**Supplementary Fig. 4. Representative photomicrographs of WT and *ATG7* KO U2OS cells stably transfected with either empty vector or cell cycle indicator PIP-FUCCI plasmid.** All cancer cell lines grew as adherent monolayer with epithelial morphology. WT Vector, wide type U2OS cells stably expressing empty vector; *ATG7* KO Vector, *ATG7* knockout U2OS cells stably expressing empty vector; WT PIP-FUCCI, wide type U2OS cells stably expressing PIP-FUCCI; *ATG7* KO PIP-FUCCI, *ATG7* knockout U2OS cells stably expressing PIP-FUCCI. Scale bars, 100 μm.

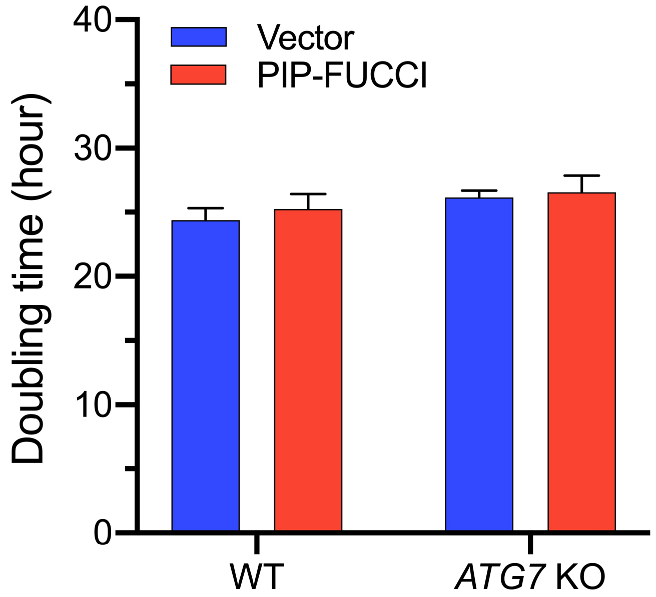

**Supplementary Fig. 5. Doubling times of WT and *ATG7* KO U2OS cells stably transfected with either empty vector or cell cycle indicator PIP-FUCCI plasmid.** Corresponding doubling times were as follows: WT Vector, 24.37 ± 0.92 hours; *ATG7* KO Vector, 26.16 ± 0.51 hours; WT PIP-FUCCI, 25.24 ± 1.17 hours; *ATG7* KO PIP-FUCCI, 26.55 ± 1.30 hours. Data were presented as "mean ± standard deviation" of three independent experiments. Statistical differences among various groups were evaluated with an independent sample *t*-test, with no statistically significant difference detected (all $P > 0.05$). WT Vector, wide type U2OS cells stably expressing empty vector; *ATG7* KO Vector, *ATG7* knockout U2OS cells stably expressing empty vector; WT PIP-FUCCI, wide type U2OS cells stably expressing PIP-FUCCI; *ATG7* KO PIP-FUCCI, *ATG7* knockout U2OS cells stably expressing PIP-FUCCI.

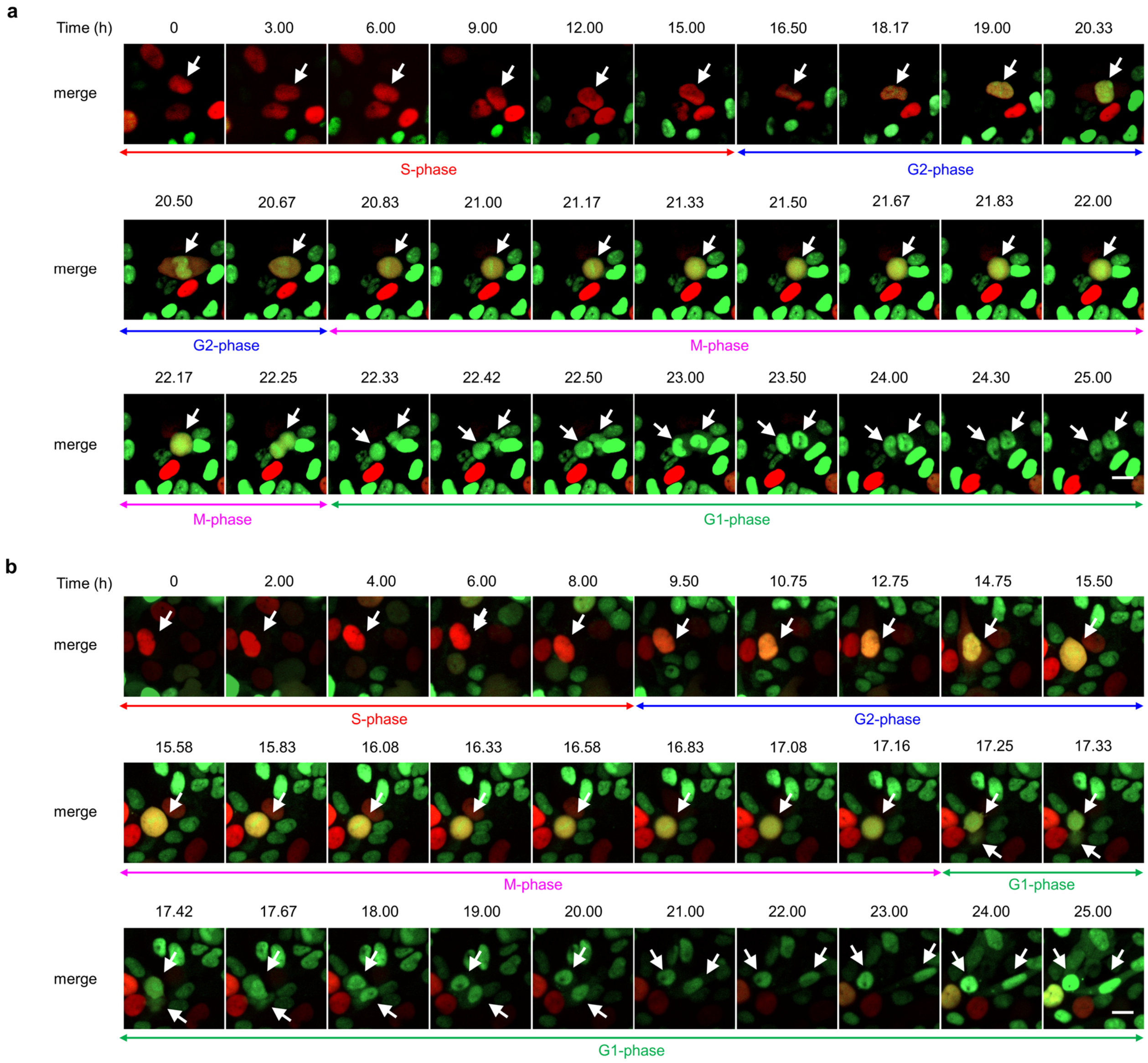

**Supplementary Fig. 6. Representative images of individually tracked WT and *ATG7* KO PIP-FUCCI cells monitored with time-lapse live cell imaging for up to 25 hours. a-b,** CLSM images showing merges of green and red fluorescence in WT (a) and *ATG7* KO (b) PIP-FUCCI cells were selected from data automatically recorded every 5 minutes for and presented at irregular intervals. Adherent cells at S-phase (with red nucleus) at time = 0 were tracked and indicated by white arrows, and their sequential cell cycle progression to adherent G2-phasc (with yellow nucleus), spherical M-phase (with yellow nucleus) and adherent Gl-phase (with green nucleus) of next cell cycle within 25 hours was clearly indicated. It is worth noting that alive cells might have tiny movements visible under CLSM. Scale bars, 20 µm.

**Supplementary Table 1. Cell cycle distribution determined with flow cytometry after 2 hour-incubation of nanoparticles**

| Cell line | Cell cycle phase | Group | | |
|---|---|---|---|---|
| | | Control (%) | DiD-LIP (%) | DiD-LNP (%) |
| WT PIP-FUCCI | G1 | 33.10 ± 1.77 | 33.18 ± 2.08 | 33.70 ± 2.36 |
| | S | 54.83 ± 2.82 | 54.64 ± 3.14 | 54.38 ± 3.37 |
| | G2 | 11.81 ± 0.93 | 11.92 ± 1.09 | 11.66 ± 0.92 |
| | M | 0.26 ± 0.19 | 0.26 ± 0.17 | 0.27 ± 0.21 |
| *ATG7* KO PIP-FUCCI | G1 | 31.30 ± 2.21 | 31.25 ± 2.15 | 31.15 ± 0.61 |
| | S | 54.22 ± 2.49 | 53.99 ± 2.37 | 55.02 ± 1.58 |
| | G2 | 13.94 ± 1.29 | 14.29 ± 0.57 | 13.27 ± 1.44 |
| | M | 0.53 ± 0.16 | 0.48 ± 0.10 | 0.56 ± 0.16 |

DiD-LIP at a dose equivalent to 5 µg/mL DOX and DiD-LNP at a dose equivalent to 6 µg/mL luciferase mRNA were added to cancer cells for 2 hours. M-phase cells were collected with mitotic shake-off as described in "Methods" before analyzing with flow cytometry. Data were presented as "mean ± standard deviation" of three independent experiments.

**Supplementary Table 2. Statistical analysis of cell cycle distribution determined with flow cytometry after 2 hour-incubation of nanoparticles**

| Cell line | Group | *P* value | | | |
|---|---|---|---|---|---|
| | | **G1-phase** | **S-phase** | **G2-phase** | **M-phase** |
| WT PIP-FUCCI | Control vs. DiD-LIP | 1.0000 | 1.0000 | 1.0000 | 1.0000 |
| | Control vs. DiD-LNP | 1.0000 | 1.0000 | 1.0000 | 1.0000 |
| | DiD-LIP vs. DiD-LNP | 1.0000 | 1.0000 | 1.0000 | 1.0000 |
| *ATG7* KO PIP-FUCCI | Control vs. DiD-LIP | 1.0000 | 1.0000 | 1.0000 | 1.0000 |
| | Control vs. DiD-LNP | 1.0000 | 1.0000 | 1.0000 | 1.0000 |
| | DiD-LIP vs. DiD-LNP | 1.0000 | 1.0000 | 0.9690 | 1.0000 |

DiD-LIP at a dose equivalent to 5 μg/mL DOX and DiD-LNP at a dose equivalent to 6 μg/mL luciferase mRNA were added to cancer cells for 2 hours. M-phase cells were collected with mitotic shake-off as described in "Methods" before analyzing with flow cytometry. Statistical analysis was conducted using ANOVA with Bonferroni correction. The level of significance was set at $P < 0.05$.

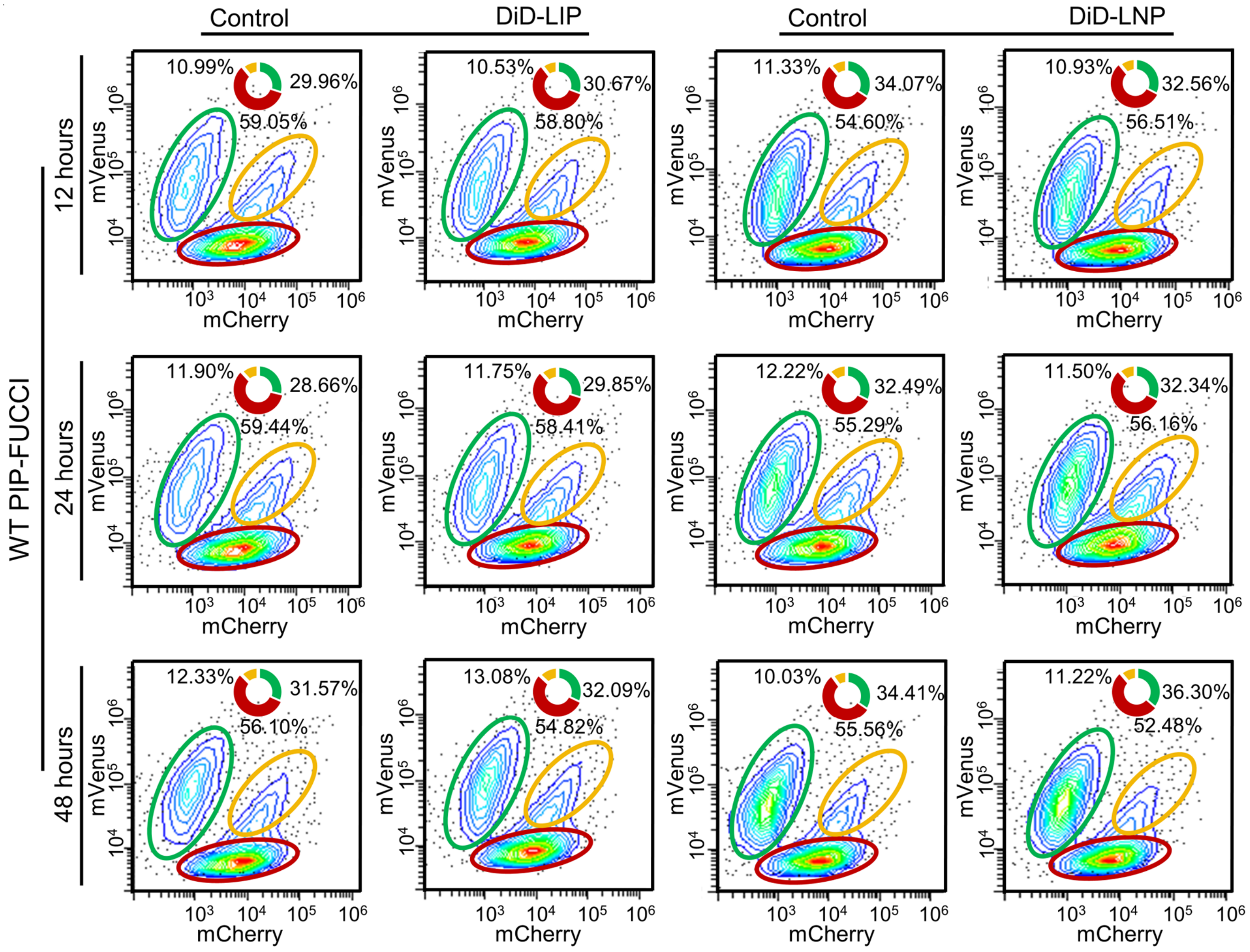

**Supplementary Fig. 7. Representative flow cytometry data on cell cycle distribution of WT PIP-FUCCI cells, with or without nanoparticle treatment at multiple time points.** DiD-LIP at a dose equivalent to 5 μg/mL DOX or DiD-LNP at a dose equivalent to 6 μg/mL luciferase mRNA were added to cancer cells for 12 hours, 24 hours and 48 hours, respectively. Afterwards, cells were routinely harvested (without specific collection of M-phase cells using mitotic shake-off) for cell cycle distribution with flow cytometry. Cells accumulated at G1-phase, S-phase and G2/M-phase were respectively circled in green, red and orange, with the corresponding percentages indicated.

**Supplementary Table 3**. **Cell cycle distribution in WT PIP-FUCCI cells after 12 hour-, 24 hour- and 48 hour-incubation of nanoparticles**

| Time | Cell cycle phase | Group | | | |
|---|---|---|---|---|---|
| | | Control (%) | DiD-LIP (%) | Control (%) | DiD-LNP (%) |
| 12 hours | G1 | 31.01 ± 0.96 | 30.43 ± 0.77 | 33.50 ± 0.71 | 33.07 ± 0.66 |
| | S | 58.22 ± 0.72 | 58.81 ± 0.65 | 55.02 ± 0.92 | 55.92 ± 0.80 |
| | G2/M | 10.77 ± 0.41 | 10.75 ± 0.22 | 11.48 ± 0.36 | 11.02 ± 0.15 |
| 24 hours | G1 | 29.22 ± 0.59 | 28.91 ± 0.90 | 32.08 ± 0.37 | 32.48 ± 0.12 |
| | S | 59.00 ± 0.40 | 59.22 ± 0.70 | 55.97 ± 0.66 | 55.57 ± 0.57 |
| | G2/M | 11.77 ± 0.24 | 11.87 ± 0.36 | 11.96 ± 0.43 | 11.95 ± 0.45 |
| 48 hours | G1 | 31.89 ± 1.58 | 33.47 ± 1.19 | 33.96 ± 1.25 | 35.11 ± 1.95 |
| | S | 55.49 ± 1.22 | 54.03 ± 0.77 | 55.03 ± 0.76 | 53.07 ± 1.70 |
| | G2/M | 12.62 ± 0.53 | 12.50 ± 0.61 | 11.01 ± 1.98 | 11.81 ± 0.52 |

DiD-LIP at a dose equivalent to 5 μg/mL DOX or DiD-LNP at a dose equivalent to 6 μg/mL luciferase mRNA were added to WT PIP-FUCCI cancer cells for 12 hours, 24 hours and 48 hours, respectively. Cells were routinely harvested (without specific collection of M-phase cells using mitotic shake-off) for cell cycle distribution with flow cytometry. Data were presented as "mean ± standard deviation" of three independent experiments. Statistical differences of the corresponding cell cycle phase between control and nanoparticle-treated groups at each time point were evaluated with an independent sample *t*-test, with no statistically significant difference detected (all $P > 0.05$).

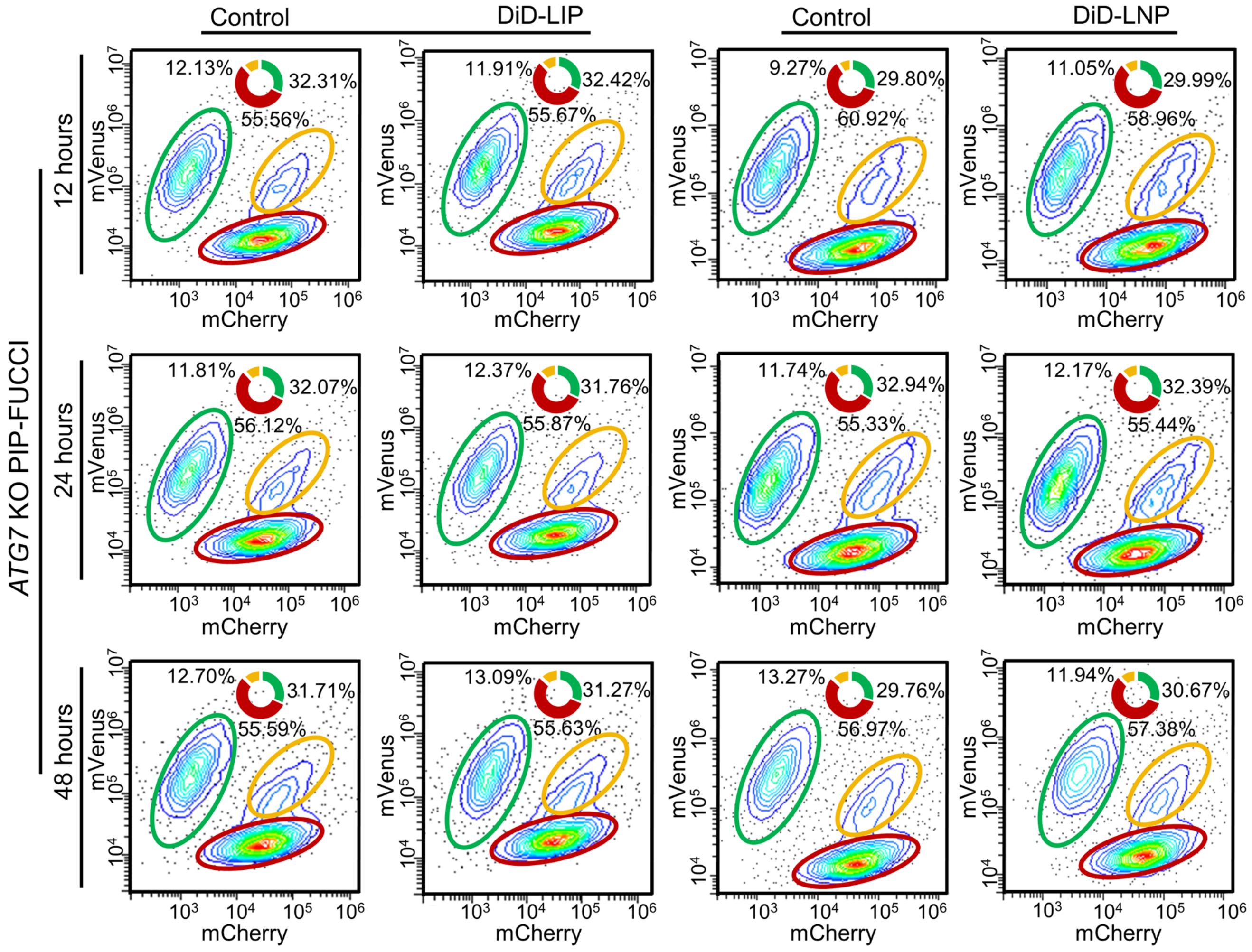

**Supplementary Fig. 8. Representative flow cytometry data on cell cycle distribution of *ATG7* KO PIP-FUCCI cells, with or without nanoparticle treatment at multiple time points.** DiD-LIP at a dose equivalent to 5 μg/mL DOX or DiD-LNP at a dose equivalent to 6 μg/mL luciferase mRNA were added to cancer cells for 12 hours, 24 hours and 48 hours, respectively. Afterwards, cells were routinely harvested (without specific collection of M-phase cells using mitotic shake-off) for cell cycle distribution with flow cytometry. Cells accumulated at G1-phase, S-phase and G2/M-phase were respectively circled in green, red and orange, with the corresponding percentages indicated.

**Supplementary Table 4**. **Cell cycle distribution in *ATG7* KO PIP-FUCCI cells after 12 hour-, 24 hour- and 48 hour-incubation of nanoparticle**

| Time | Cell cycle phase | Group | | | |
|---|---|---|---|---|---|
| | | Control (%) | DiD-LIP (%) | Control (%) | DiD-LNP (%) |
| 12 hours | G1 | 31.70 ± 1.23 | 32.15 ± 0.99 | 30.62 ± 0.72 | 31.96 ± 2.60 |
| | S | 55.85 ± 0.76 | 56.24 ± 0.80 | 59.98 ± 0.82 | 58.18 ± 1.38 |
| | G2/M | 12.45 ± 0.48 | 11.62 ± 0.42 | 9.40 ± 0.13 | 9.86 ± 1.28 |
| 24 hours | G1 | 32.29 ± 0.28 | 32.37 ± 0.53 | 32.36 ± 0.59 | 32.56 ± 0.52 |
| | S | 56.33 ± 0.52 | 55.77 ± 0.76 | 55.94 ± 0.56 | 55.53 ± 0.96 |
| | G2/M | 11.37 ± 0.48 | 11.86 ± 0.89 | 11.71 ± 0.12 | 11.91 ± 0.51 |
| 48 hours | G1 | 31.09 ± 0.58 | 31.90 ± 0.62 | 30.81 ± 1.01 | 31.23 ± 0.52 |
| | S | 56.82 ± 1.11 | 55.56 ± 0.11 | 56.49 ± 0.62 | 56.62 ± 0.73 |
| | G2/M | 12.09 ± 0.54 | 12.54 ± 0.52 | 12.70 ± 0.49 | 12.15 ± 0.22 |

DiD-LIP at a dose equivalent to 5 µg/mL DOX or DiD-LNP at a dose equivalent to 6 µg/mL luciferase mRNA were added to *ATG7* KO PIP-FUCCI cells for 12 hours, 24 hours and 48 hours, respectively. Cells were routinely harvested (without specific collection of M-phase cells using mitotic shake-off) for cell cycle distribution with flow cytometry. Data were presented as "mean ± standard deviation" of three independent experiments. Statistical differences of the corresponding cell cycle phase between control and nanoparticle-treated groups at each time point were evaluated with an independent sample *t*-test, with no statistically significant difference detected (all $P > 0.05$).

**Supplementary Table 5. Autophagic vesicle (AV) numbers in WT and *ATG7* KO PIP-FUCCI cells and subsequent statistical analysis**

| Group | WT PIP-FUCCI | *ATG7* KO PIP-FUCCI |
|---|---|---|
| Control (AV number per cell) | 3.60 ± 2.32 | 0.30 ± 0.48 |
| DiD-LIP (AV number per cell) | 3.30 ± 1.95 | 0.20 ± 0.63 |
| DiD-LNP (AV number per cell) | 3.40 ± 1.51 | 0.10 ± 0.32 |
| Control vs. DiD-LIP (*P* value 1) | 0.8182 | 0.3566 |
| Control vs. DiD-LNP (*P* value 2) | 0.9695 | 0.3566 |
| DiD-LIP vs. DiD-LNP (*P* value 3) | 0.8990 | 0.6600 |

AVs were examined with transmission electron microscopy. AV numbers were presented as "mean ± standard deviation" (n = 10). Statistical analysis of AV numbers among various groups was conducted using Mann-Whitney U test, where *P* value 1 represented differences between Control and DiD-LIP groups, *P* value 2 represented differences between Control and DiD-LNP groups, *P* value 3 represented differences between DiD-LIP and DiD-LNP groups. The level of significance was set at $P < 0.05$.

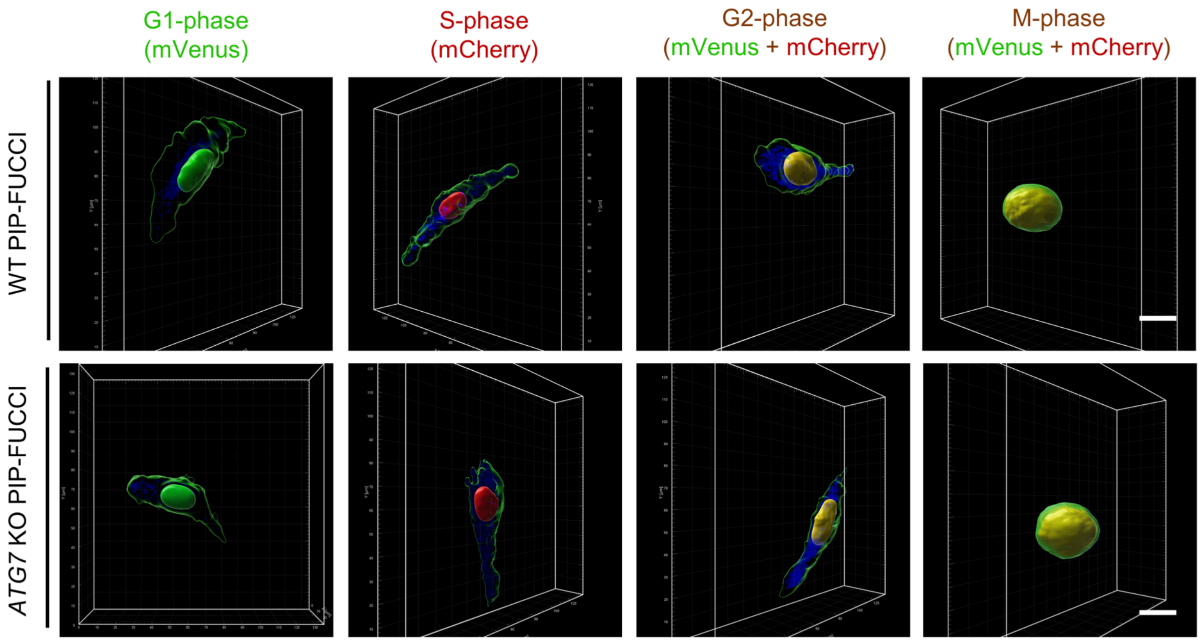

**Supplementary Fig. 9. Representative screenshots of three-dimensional reconstruction images for determining cell cycle- and autophagy-associated cellular uptake of DiD-labelled liposomes (DiD-LIP) using the multi-functional platform.** DiD-LIP (blue fluorescence) was administered to cancer cells for 2 hours at a dose equivalent to 5 μg/mL DOX. Adherent G1-phase cells, mVenus positive, green nucleus; adherent S-phase cells, mCherry positive, red nucleus; adherent G2-phase cells and detached M-phase cells collected with mitotic shake-off, mVenus and mCherry double-positive, yellow nucleus. WT PIP-FUCCI, wide type U2OS cells stably expressing PIP-FUCCI; *ATG7* KO PIP-FUCCI, *ATG7* knockout U2OS cells stably expressing PIP-FUCCI. Scale bars, 20 μm.

**Supplementary Table 6**. **Cell area, cell volume, TFI and MFI of DiD-LIP acquired based on the multi-functional platform**

| Cell line | Cell cycle phase | Cell area (pixel) | Cell volume (μm$^3$) | TFI (a.u.) | | MFI (a.u.) | |
|---|---|---|---|---|---|---|---|
| | | | | 2D | 3D | 2D | 3D |
| WT PIP-FUCCI | G1 | 5.058 ± 0.185 | 3.725 ± 0.211 | 6.089 ± 0.214 | 8.966 ± 0.214 | 1.031 ± 0.097 | 5.241 ± 0.132 |
| | S | 5.150 ± 0.138 | 3.821 ± 0.136 | 6.189 ± 0.139 | 9.067 ± 0.111 | 1.039 ± 0.118 | 5.246 ± 0.126 |
| | G2 | 5.293 ± 0.204 | 3.981 ± 0.173 | 6.368 ± 0.196 | 9.268 ± 0.198 | 1.074 ± 0.089 | 5.287 ± 0.101 |
| | M | 5.099 ± 0.182 | 3.984 ± 0.290 | 5.690 ± 0.204 | 8.863 ± 0.269 | 0.591 ± 0.182 | 4.879 ± 0.176 |
| *ATG7* KO PIP-FUCCI | G1 | 5.210 ± 0.128 | 3.830 ± 0.144 | 6.258 ± 0.142 | 9.085 ± 0.158 | 1.047 ± 0.131 | 5.255 ± 0.127 |
| | S | 5.256 ± 0.131 | 3.932 ± 0.148 | 6.297 ± 0.156 | 9.234 ± 0.139 | 1.041 ± 0.116 | 5.303 ± 0.096 |
| | G2 | 5.386 ± 0.153 | 4.079 ± 0.186 | 6.439 ± 0.163 | 9.349 ± 0.136 | 1.053 ± 0.105 | 5.319 ± 0.114 |
| | M | 5.064 ± 0.106 | 4.012 ± 0.178 | 5.772 ± 0.178 | 8.959 ± 0.168 | 0.708 ± 0.214 | 4.947 ± 0.226 |

Data were first analyzed after log$_{10}$ transformation and presented as "mean ± standard deviation" (n = 25-30). TFI, Total fluorescence intensity; MFI, Mean fluorescence intensity; 2D, Two-dimensional level; 3D, Three-dimensional level.

**Supplementary Table 7**. **Statistical analysis of cell area, cell volume, TFI and MFI of DiD-LIP among four different cell cycle phases**

| **Cell line** | **Cell cycle phase** | ***P* value** | | | | | |
|---|---|---|---|---|---|---|---|
| | | **Cell area** | **Cell volume** | **TFI (2D)** | **TFI (3D)** | **MFI (2D)** | **MFI (3D)** |
| WT PIP-FUCCI | G1 vs. S | 0.2890 | 0.4600 | 0.2650 | 0.3530 | 1.0000 | 1.0000 |
| | G1 vs. G2 | < 0.0001 | < 0.0001 | < 0.0001 | < 0.0001 | 1.0000 | 1.0000 |
| | G1 vs. M | 1.0000 | < 0.0001 | < 0.0001 | 0.3510 | < 0.0001 | < 0.0001 |
| | S vs. G2 | 0.0140 | 0.0230 | 0.0030 | 0.0015 | 1.0000 | 1.0000 |
| | S vs. M | 1.0000 | 0.0210 | < 0.0001 | 0.0014 | < 0.0001 | < 0.0001 |
| | G2 vs. M | < 0.0001 | 1.0000 | < 0.0001 | < 0.0001 | < 0.0001 | < 0.0001 |
| *ATG7* KO PIP-FUCCI | G1 vs. S | 1.0000 | 0.1201 | 1.0000 | 0.0015 | 1.0000 | 1.0000 |
| | G1 vs. G2 | < 0.0001 | < 0.0001 | 0.0002 | < 0.0001 | 1.0000 | 0.5492 |
| | G1 vs. M | 0.0004 | 0.0005 | < 0.0001 | 0.0153 | < 0.0001 | < 0.0001 |
| | S vs. G2 | 0.0017 | 0.0057 | 0.0060 | 0.0267 | 1.0000 | 1.0000 |
| | S vs. M | < 0.0001 | 0.4779 | < 0.0001 | < 0.0001 | < 0.0001 | < 0.0001 |
| | G2 vs. M | < 0.0001 | 0.8164 | < 0.0001 | < 0.0001 | < 0.0001 | < 0.0001 |

Statistical analysis among four different cell cycle phases was conducted using ANOVA with Bonferroni correction. The level of significance was set at $P < 0.05$. TFI, Total fluorescence intensity; MFI, Mean fluorescence intensity; 2D, Two-dimensional level; 3D, Three-dimensional level.

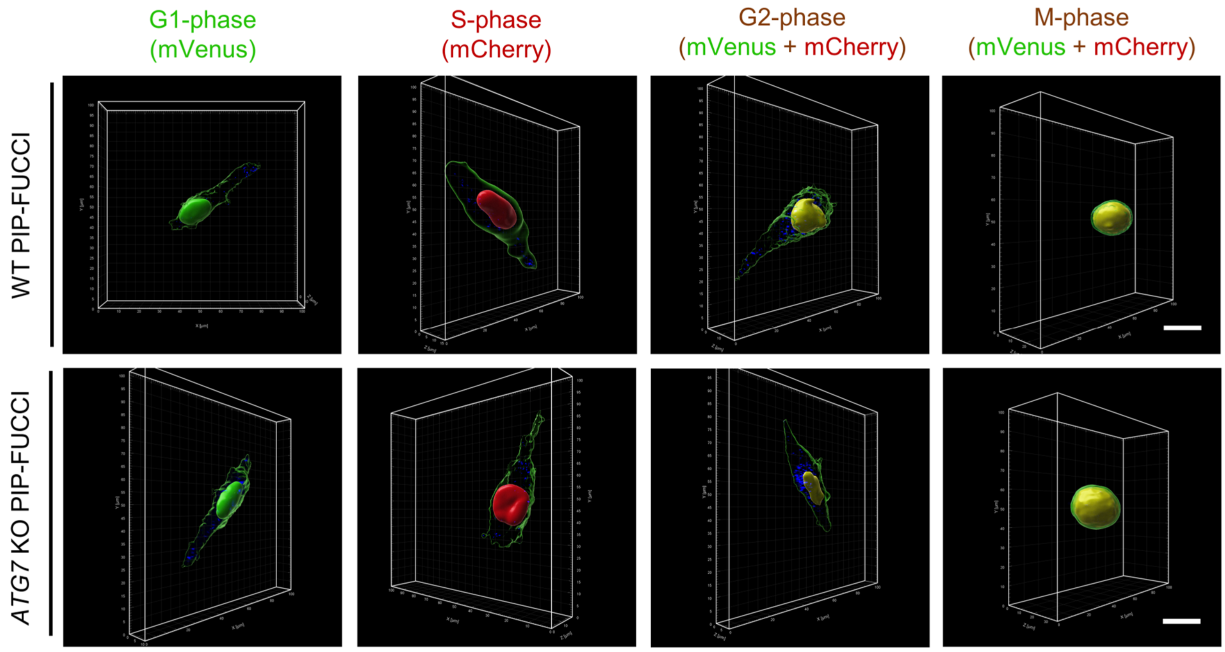

**Supplementary Fig. 10. Representative screenshots of three-dimensional reconstruction images for determining cell cycle- and autophagy-associated cellular uptake of DiD-labelled lipid nanoparticles (DiD-LNP) using the multi-functional platform.** DiD-LNP (blue fluorescence) was administered to cancer cells for 2 hours at a dose equivalent to 6 μg/mL luciferase mRNA. Adherent G1-phase cells, mVenus positive, green nucleus; adherent S-phase cells, mCherry positive, red nucleus; adherent G2-phase cells and detached M-phase cells collected with mitotic shake-off, mVenus and mCherry double-positive, yellow nucleus. WT PIP-FUCCI, wide type U2OS cells stably expressing PIP-FUCCI; *ATG7* KO PIP-FUCCI, *ATG7* knockout U2OS cells stably expressing PIP-FUCCI. Scale bars, 20 μm.

**Supplementary Table 8**. **Cell area, cell volume, TFI and MFI of DiD-LNP acquired based on the multi-functional platform**

| Cell line | Cell cycle phase | Cell area (pixel) | Cell volume (μm³) | TFI (a.u.) | | MFI (a.u.) | |
|---|---|---|---|---|---|---|---|
| | | | | 2D | 3D | 2D | 3D |
| WT PIP-FUCCI | G1 | 5.184 ± 0.196 | 3.934 ± 0.167 | 5.754 ± 0.322 | 8.617 ± 0.237 | 0.570 ± 0.280 | 4.683 ± 0.246 |
| | S | 5.181 ± 0.128 | 3.806 ± 0.149 | 5.845 ± 0.306 | 8.715 ± 0.268 | 0.665 ± 0.238 | 4.909 ± 0.229 |
| | G2 | 5.231 ± 0.140 | 3.964 ± 0.175 | 6.147 ± 0.143 | 9.118 ± 0.206 | 0.916 ± 0.147 | 5.154 ± 0.154 |
| | M | 5.044 ± 0.180 | 4.074 ± 0.272 | 4.979 ± 0.539 | 8.221 ± 0.481 | -0.064 ± 0.479 | 4.148 ± 0.350 |
| *ATG7* KO PIP-FUCCI | G1 | 5.249 ± 0.135 | 4.059 ± 0.144 | 5.757 ± 0.294 | 8.751 ± 0.243 | 0.508 ± 0.284 | 4.692 ± 0.265 |
| | S | 5.353 ± 0.146 | 4.059 ± 0.172 | 5.762 ± 0.230 | 8.700 ± 0.238 | 0.410 ± 0.253 | 4.641 ± 0.243 |
| | G2 | 5.429 ± 0.190 | 4.169 ± 0.188 | 6.011 ± 0.366 | 9.050 ± 0.333 | 0.581 ± 0.306 | 4.882 ± 0.325 |
| | M | 5.233 ± 0.120 | 4.357 ± 0.157 | 5.358 ± 0.245 | 8.573 ± 0.204 | 0.125 ± 0.232 | 4.215 ± 0.197 |

Data were first analyzed after $\log_{10}$ transformation and presented as "mean ± standard deviation" (n = 25-30). TFI, Total fluorescence intensity; MFI, Mean fluorescence intensity; 2D, Two-dimensional level; 3D, Three-dimensional level.

**Supplementary Table 9. Statistical analysis of cell area, cell volume, TFI and MFI of DiD-LNP among four different cell cycle phases**

| **Cell line** | **Cell cycle phase** | ***P* value** | | | | | |
|---|---|---|---|---|---|---|---|
| | | **Cell area** | **Cell volume** | **TFI (2D)** | **TFI (3D)** | **MFI (2D)** | **MFI (3D)** |
| WT PIP-FUCCI | G1 vs. S | 1.0000 | 0.0930 | 1.0000 | 1.0000 | 1.0000 | 0.0070 |
| | G1 vs. G2 | 1.0000 | 1.0000 | 0.0006 | < 0.0001 | 0.0005 | < 0.0001 |
| | G1 vs. M | 0.0080 | 0.0440 | < 0.0001 | < 0.0001 | < 0.0001 | < 0.0001 |
| | S vs. G2 | 1.0000 | 0.0250 | 0.0170 | < 0.0001 | 0.0260 | 0.0040 |
| | S vs. M | 0.0120 | < 0.0001 | < 0.0001 | < 0.0001 | < 0.0001 | < 0.0001 |
| | G2 vs. M | < 0.0001 | 0.2470 | < 0.0001 | < 0.0001 | < 0.0001 | < 0.0001 |
| *ATG7* KO PIP-FUCCI | G1 vs. S | 0.0570 | 1.0000 | 1.0000 | 1.0000 | 1.0000 | 1.0000 |
| | G1 vs. G2 | < 0.0001 | 0.1020 | 0.0090 | < 0.0001 | 1.0000 | 0.0530 |
| | G1 vs. M | 1.0000 | < 0.0001 | < 0.0001 | 0.0690 | < 0.0001 | < 0.0001 |
| | S vs. G2 | 0.3560 | 0.0920 | 0.0098 | < 0.0001 | 0.1140 | 0.0050 |
| | S vs. M | 0.0180 | < 0.0001 | < 0.0001 | 0.3910 | 0.0008 | < 0.0001 |
| | G2 vs. M | < 0.0001 | < 0.0001 | < 0.0001 | < 0.0001 | < 0.0001 | < 0.0001 |

Statistical analysis among four different cell cycle phases was conducted using ANOVA with Bonferroni correction. The level of significance was set at $P < 0.05$. TFI, Total fluorescence intensity; MFI, Mean fluorescence intensity; 2D, Two-dimensional level; 3D, Three-dimensional level.

**Supplementary Table 10. Statistical analysis on correlation coefficients among TFI and MFI in groups treated with different nanoparticles**

| Cell line | Correlation coefficient | DiD-LIP vs. DiD-LNP |
|---|---|---|
| WT PIP-FUCCI | TFI (2D) and TFI (3D) | $P = 0.0441$ |
| | MFI (2D) and MFI (3D) | $P < 0.0001$ |
| | MFI (2D) and TFI (2D) | $P < 0.0001$ |
| | MFI (3D) and TFI (3D) | $P < 0.0001$ |
| *ATG7* KO PIP-FUCCI | TFI (2D) and TFI (3D) | $P = 0.0860$ |
| | MFI (2D) and MFI (3D) | $P < 0.0001$ |
| | MFI (2D) and TFI (2D) | $P = 0.0023$ |
| | MFI (3D) and TFI (3D) | $P = 0.0007$ |

DiD-LIP at a dose equivalent to 5 μg/mL DOX and DiD-LNP at a dose equivalent to 6 μg/mL luciferase mRNA were added to cancer cells for 2 hours. Statistical analysis was conducted using z-test on Fisher z-transformed correlation coefficients. The level of significance was set at $P < 0.05$.

**Supplementary Table 11. Statistical analysis on correlation coefficients among TFI and MFI in WT and *ATG7* KO PIP-FUCCI cells**

| Nanoparticle type | Correlation coefficient | WT PIP-FUCCI vs. *ATG7* KO PIP-FUCCI |
|---|---|---|
| DiD-LIP | TFI (2D) and TFI (3D) | *P* = 0.1841 |
| | MFI (2D) and MFI (3D) | *P* = 0.0795 |
| | MFI (2D) and TFI (2D) | *P* = 0.8577 |
| | MFI (3D) and TFI (3D) | *P* = 0.5989 |
| DiD-LNP | TFI (2D) and TFI (3D) | *P* = 0.1178 |
| | MFI (2D) and MFI (3D) | *P* = 0.1974 |
| | MFI (2D) and TFI (2D) | *P* = 0.0275 |
| | MFI (3D) and TFI (3D) | *P* = 0.0516 |

DiD-LIP at a dose equivalent to 5 µg/mL DOX and DiD-LNP at a dose equivalent to 6 µg/mL luciferase mRNA were added to cancer cells for 2 hours. Statistical analysis was conducted using z-test on Fisher z-transformed correlation coefficients. The level of significance was set at $P < 0.05$.

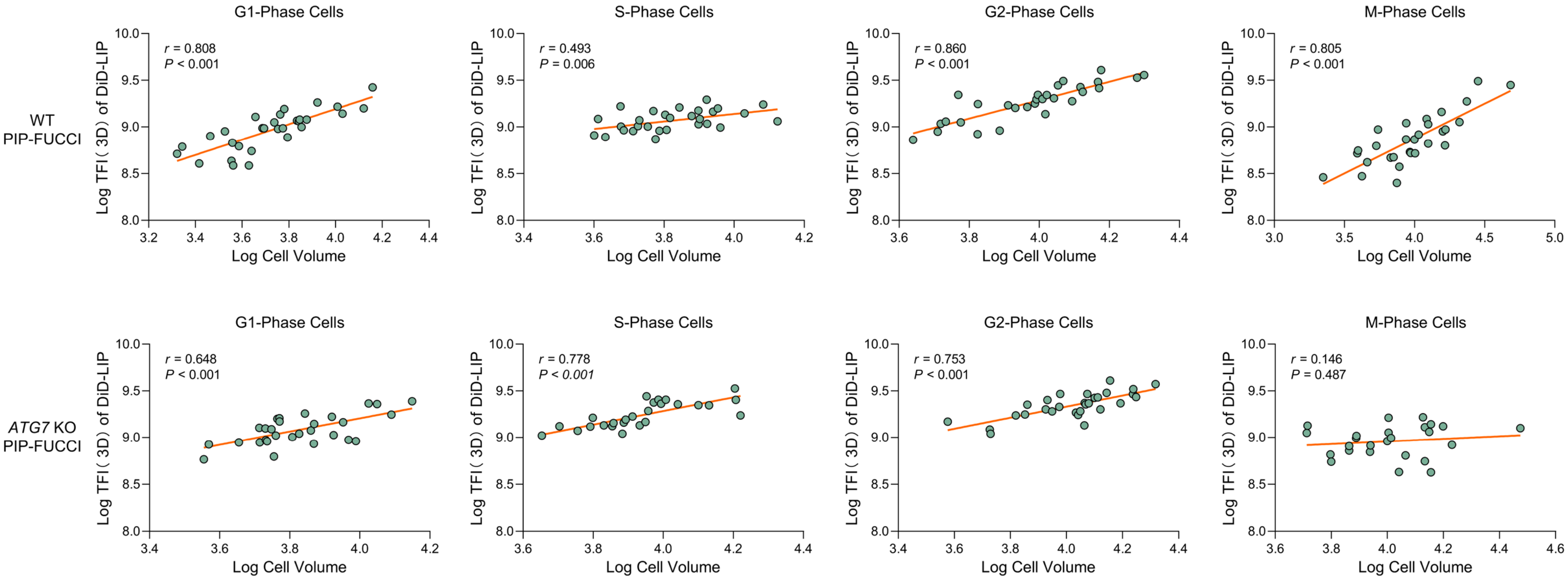

**Supplementary Fig. 11**. **Pearson correlation coefficients of cell volume and TFI (3D) of DiD-LIP at each cell cycle phase in WT and *ATG7* KO PIP-FUCCI cells.** The nonparametric test, Pearson correlation coefficient was used to evaluate the correlation of $\log_{10}$ transformed DiD-LIP TFI (3D) and $\log_{10}$ transformed cell volume. Level of significance was set at $P < 0.05$, and correlation strength was determined using correlation coefficient ($r$).

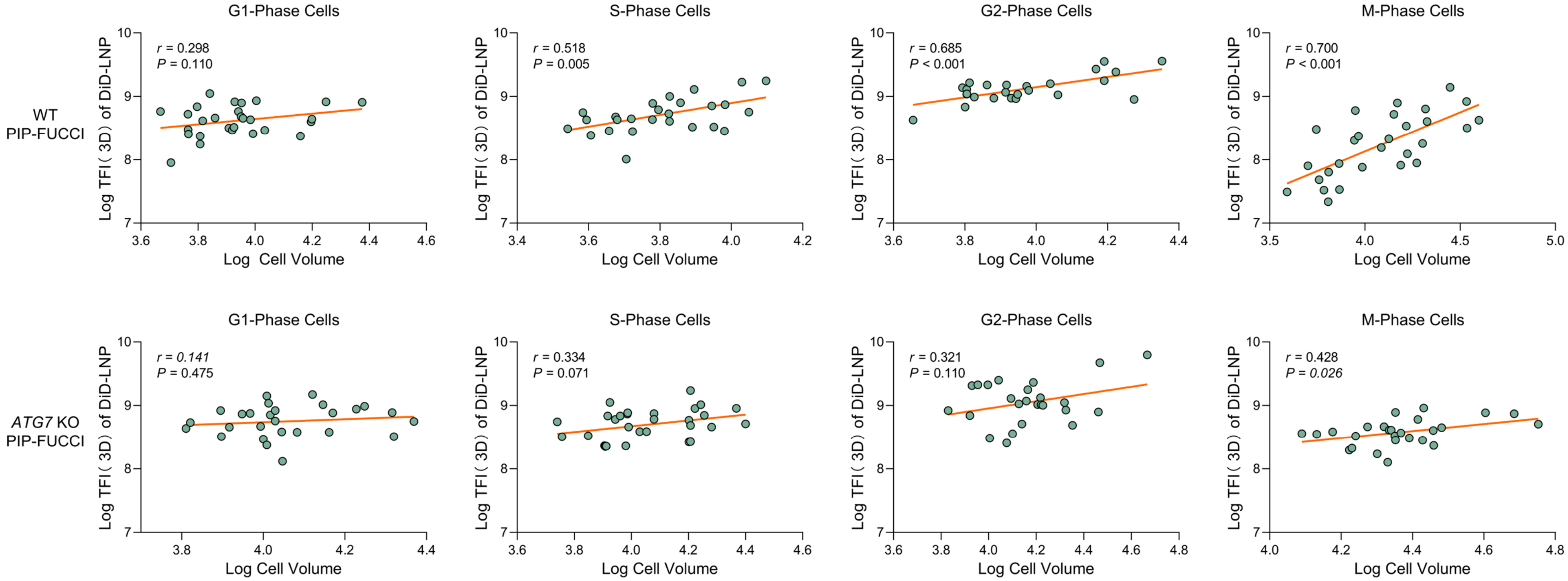

**Supplementary Fig. 12**. **Pearson correlation coefficients of cell volume and TFI (3D) of DiD-LNP at each cell cycle phase in WT and *ATG7* KO PIP-FUCCI cells.** The nonparametric test, Pearson correlation coefficient was used to evaluate the correlation of $\log_{10}$ transformed DiD-LIP TFI (3D) and $\log_{10}$ transformed cell volume. Level of significance was set at $P < 0.05$, and correlation strength was determined using correlation coefficient ($r$).
